# Fossil Freshwater Fishes from the Pliocene of northern Colombia and the Palaeogeography of northern South America

**DOI:** 10.1101/2024.12.18.629252

**Authors:** Gustavo A. Ballen, Camilo Montes, Jorge D. Carrillo-Briceño, Sergio Bogan, Sandra Reinales, Mario C. C. de Pinna, Carlos Jaramillo

## Abstract

Freshwater fishes from northern Colombia are reported from the Sincelejo and Ware formations, of Pliocene age. A total of ten taxa have been identified comprising two orders, five families, and nine genera. Characters from dental morphology, fin spines, and cranial bones are provided as taxonomic tools for the study of fossil fishes. All of the taxa are members of groups currently restricted to drainages east of the Andes, suggesting that physical drainage connection was still present by the Pliocene between the Amazon-Orinoco and trans-Andean drainages such as the Magdalena-Cauca, or that these groups persisted in the trans-Andean region at least until Pliocene times. The genera *Hemidoras*, *Serrasalmus*, and *Trachelyopterichthys* are new records for the fossil fish fauna of South America. The genus *Zungaro* represents a new record for the trans-Andean region, whereas the genus *Platysilurus* is for the first time in Colombia. Most of these occurrences also represent the youngest occurrences in the fossil record. Literature records are reidentified as *Pygocentrus* from the La Venta fauna and the earily Oligocene in Peru. These assemblages suggest that the Sincelejo and Ware formations were deposited in rivers of large size that were part of a large drainage network connected to the Amazon-Orinoco despite being currently located west of the Andes. These findings suggest that the Cordillera Oriental and the Merida Andes were not dividing yet the drainage network in northern South America by the middle to late Miocene.

## 2 Introduction

Freshwater fishes in the Neotropics are one of the most speciose vertebrate groups in the world with thousands of species occupying a variety of habitats ranging from sea level to ¿ 4000 masl in the Andes (Albert et al., 2011). They are ecologically diverse and play a key role in aquatic habitats in the continent, as well as food resource for human populations (Carolsfeld et al., 2003). This diversity is coupled with a proportional morphological variation and large body size variance spanning several orders of magnitude from miniature catfishes (Schaefer et al., 1989) to the giant bonytongue *Arapaima* (Castello, 2008). This high diversity has been the product of a large history of evolutionary change, diversification, and adaptation in South America (Lundberg, 1997; Lundberg et al., 1998; Albert et al., 2006, 2011).

The Andean orogeny has been a key geologic process affecting the biota of South America (Antonelli et al., 2009; Hoorn et al., 2010). Its effect on the diversification of different living groups has been shown along its entire area of influence from the limit between Chile and Argentina up to Venezuela via creation of new environments, vicariance of pre-existing wider geographic distributions, and the consequent drainage separation in the case of aquatic organisms. The uplift timing of the Andes, and especially that of the northern Andes is still highly controversial. Some authors suggest that the Cordillera Oriental in Colombia, and the Mérida Andes and Coastal Cordillera in Venezuela were positive by middle to late Miocene and the Perijá Range and the Mérida Andes during the late Miocene (Diaz de Gamero, 1996; Lundberg et al., 1998, and references therein). In contrast, some authors suggest a more complex tectonic history for the Andes of northern South America (NSA), arguing that the Cordillera Oriental in Colombia uplifted in pulses, with some areas being positive as early as late Palaeocene (e.g., Santander Massif; Bayona et al., 2013) and with other areas uplifting from middle Eocene to late Miocene (e.g., Perijá Range and Cordillera Oriental; Ayala et al., 2012; Bayona et al., 2010, 2013; Caballero et al., 2010; Ochoa et al., 2012). Both scenarios would have different consequences for the freshwater fish faunas (both extant and extinct), and therefore, they could be used to better understand the evolution of the northern Andes. Palaeoenvironmental reconstructions suggest that the Peninsula was a tropical forest with presence of either middle to large rivers or a deltaic area, aquatic vertebrates such as crocodylians, turtles, freshwater fishes, as well as terrestrial and amphibian mammals (Aguilera et al., 2013; Forasiepi et al., 2014; Cadena and Jaramillo, 2015b,a; Amson et al., 2016; Suarez et al., 2016; Moreno-Bernal et al., 2016; Carrillo et al., 2018; Ballen et al., 2021, 2022). Such palaeobiotic assemblage allows to explore past drainage connections with major drainages of NSA because of its intermediate location between the Magdalena and Maracaibo drainages, as well as being complementary to other fossil faunas of similar age (i.e., San Gregorio formation in Venezuela). The main goal of this study is to study the freshwater fish assemblage of the Sincelejo and Ware formations in the departments of Sucre and Guajira during the Pliocene and to explore its relevance for palaeogeographic models using past and present geographic distribution of the taxa recovered.

## 3 Materials and Methods

### 3.1 Abbreviations and comparative material

Preserved specimens of extant species of the orders Characiformes and Siluriformes were examined in scientific collections (Supplementary Material S1). A total of five fossil samples were examined from the Sincelejo formation, and 116 from the Ware formation. Institutional abbreviations include: Academy of Natural Sciences of Drexel University, Philadelphia, US (ANSP), Instituto Alexander von Humboldt, Villa de Leyva, Colombia (IAvH), Museu de Zoologia da Universidade de São Paulo, São Paulo, Brazil (MZUSP), Mapuka Museum of Universidad del Norte, Barranquilla, Colombia (MUN). Premaxilla and dentary are abbreviated PM and D respectively. Specimens identified in the literature as either *Hydrolycus* or *Rhaphiodon* by Cione and Casciotta (1997) are herein restricted to the genus *Hydrolycus* instead. Unpublished records of the genus *Oxydoras* from the Ituzaingo formation by one of us (S. Bogan) are herein included for the faunal similarity analysis.

### 3.2 Anatomical terminology

Siluriform osteological terminology follows Lundberg and McDade (1986) with refinements by Slobodian and Pastana (2018). Siluriform appendicular terminology follows Ballen and de Pinna (2022, 2024). Cynodontid cranial nomenclature follows Toledo-Piza (2000) and Ballen et al. (2022). Serrasalmid tooth nomenclature and position follows Cione et al. (2009) with modifications by Ballen et al. (2021).

### 3.3 Data analysis

#### 3.3.1 Phylogenetic inference

We analyzed a combined molecular and morphological dataset with 84 terminals representing 24 genera of the Pimelodidae and two genera of the Pseudopimelodidae and the Heptapteridae respectively, while the fossil taxon set was composed of the three known extinct species of *Phractocephalus* as well as the specimens herein studied, for a total of four fossil terminals. DNA sequences for the molecular partitions were taken from Lundberg et al. (2011), who included the single-copy nuclear recombination activating genes (*rag1* and *rag2*), a mitochondrial region comprising the *12S* rRNA, tRNA-val and *16S* rRNA genes, and the cytochrome-b *cytb* gene (including Threonine tRNA and partial Proline tRNA regions), for a total of four molecular partitions. All available sequences were retrieved from GenBank and aligned with MAFFT v.7.271 (Katoh and Standley, 2013) using the G-INS-i algorithm with 1000 iterations except for *rag1* where we used the L-INS-i algorithm due to the unequal size of sequences in this partition. Minor manual adjustments were carried out for the *rag1* and *rag2* alignments. Additionally, we used Gblocks v.0.91 (Castresana, 2000; Talavera and Castresana, 2007) for reproducible exclusion of ambiguous and hypervariable positions of the *12S* alignment with the following settings: -t=d -b=a -d=y; a total of 84 (4%) positions were removed from the original alignment. Accession numbers are available below; missing data were coded as “?”. A total dataset with 84 terminals and 7578 characters was assembled for analysis.

Phylogenetic analysis was performed using Bayesian Inference as implemented in MrBayes v.3.2.6 (Ronquist et al., 2012). Substitution models for each partition were selected using JModeltest 2.1.10 v.20160303 (Darriba et al., 2012) and PhyML 3.0 v.20131022 (Guindon et al., 2010) using the Bayesian Information Criterion (BIC) following Darriba et al. (2012). The best-fit substitution models were GTR+I+Γ (*rag1*); K80+I+Γ (*rag2*); GTR+I+Γ (*12S*), and HKY+I+Γ (*cytb*). Parameters other than topology were unlinked across partitions. Two runs with eight independent Markov chains were run in parallel for 2.000.000 generations and sampling every 2.000 generations. Convergence of runs was determined based on the average standard deviation of the split frequencies (ASDSF) < 0.01, and the effective sample size (ESS), calculated using Tracer v1.6.0 (Drummond et al., 2012), that was > 300 for all parameters. Additionally, the potential scale reduction factor (PSRF) approached 1.0, suggesting convergence in the estimation of the posterior probabilities of nodes and branch length parameters. The posterior density graphs of the two independent runs were also examined in Tracer, and not visual differences were found between them. The 25% of trees were discarded as burn-in and a 50% majority-rule tree and posterior probabilities (PP) for node support were calculated using the remaining trees. Detailed description of the data and implementation of the analysis can be found in the Supplementary Material S2.

#### 3.3.2 Faunal similarity

Fossil freshwater fish assemblages are known from Colombia and Venezuela to Argentina (Lundberg et al., 2010). An analysis of assemblage composition (dis)similarity was carried out using data compiled from literature data as well as new records (Agnolin and Bogan, 2020; Aguilera et al., 2008; Aguilera and Lundberg, 2010; Aguilera et al., 2013; Antoine et al., 2016; Azpelicueta and Cione, 2016; Ballen and Moreno-Bernal, 2019; Ballen et al., 2021, 2022; Bogan et al., 2012; Bogan and Agnolín, 2019; Bogan and Agnolin, 2020a, 2021, 2020b; Carrillo-Briceño et al., 2015, 2021b,a; Carrillo-Briceño et al., 2023; Cione and Casciotta, 1997; Cione et al., 2000, 2009; Cione and Azpelicueta, 2013; Dahdul, 2004; Deynat and Brito, 1994; Lundberg and McDade, 1986; Lundberg et al., 1988; Lundberg and Chernoff, 1992; Lundberg, 1997; Lundberg and Aguilera, 2003; Lundberg, 2005; Lundberg et al., 2010; Monsch, 1998; Núñez-Flores et al., 2017; Priem, 1911; Reis, 1998; Richter, 1989; Rincón et al., 2016; Sabaj Pérez et al., 2007; Sánchez-Villagra and Aguilera, 2006; Schwarzhans et al., 2022; Tejada-Lara et al., 2015). An initial set of Miocene fossil fish faunas including both marine and freshwater componentswas included. This initial assemblage was then reduced to a set where the number of taxa was equal or larger than the one herein described. Although the original dataset includes fossil assemblages of strictly freshwater, strictly marine, or mixed origin, only those with freshwater components were included in the final analysis. The final dataset includes the Acre, Contamana, La Venta, Fitzcarrald, Ituzaingó, Makaraipao, Urumaco, and Ware fossil assemblages along with their fossil fish occurrences. The overall results using all the data or just the subset described above did not change the conclusions of the similarity analysis. Faunal similarity and resampling nodal support values were measured using the approach described in Ballen et al. (2021) using the binary method with average distance as implemented in the pvclust package v.2.2-0 (Suzuki et al., 2019) and vegan v.2.5-5 (Oksanen et al., 2019) in R v.3.4.4 R Core Development Team (2018). All the analyses were run on the Brycon server at IBB/UNESP Botucatu.

### 3.4 Geological Setting

The fossil localities from the Ware and Sincelejo formation herein studied belong to two geologic regions of northern Colombia respectively: the Cocinetas sedimentary basin and the San Jacinto tectonic belt.

#### 3.4.1 The Cocinetas sedimentary basin

The Cocinetas sedimentary basin is composed of units recording the environmental dynamics during the Eocene to Pliocene timespan (Renz, 1960; Rollins, 1965). It is bounded by the Macuira Fault to the northeast, the Cuisa Fault to the southwest, and the Serranía de Jarara to the northwest. The basin genesis has been associated to regional tectonics involving the migration of tectonic blocks and the subsequent formation of pull-apart basins along the northern margin of South America (Moreno et al., 2015). It is located to the east of the Guajira Peninsula in northern Colombia, along the international border with Venezuela.

The Ware formation is a continental succession of medium to coarse sandstone intercalated with fine levels of mudstone and some prominent levels of conglomerate and conglomeratic sandstone bounded by an uncomformity with the Castilletes formation to the base and a coquina of regional extent to the top; the preserved succession is generally thin (ca. 25 m) but can be followed regionally.. The formation preserves a rich vertebrate assemblage (Ballen et al., 2022; Amson et al., 2016; Moreno-Bernal et al., 2016; Carrillo et al., 2018; Forasiepi et al., 2014; Carrillo-Briceño et al., 2019) (Table 1; Figure 3), that has been dated as Pliocene based on radiometric data from Sr isotopes (Hendy et al., 2015). Aguilera et al. (2013) had reported a few Catfishes of the families Doradidae and Pimelodidaeand our study adds many specimens to its freshwater fish record.

**Table 1:** STRI localities from the Ware formation referenced in different sources since 2013. Several localities refer to the same level and spatial point, being therefore synonyms.

| Locality | Strat (m) | Taxon | Source | Comments |
| --- | --- | --- | --- | --- |
| 290045 | NA | NA | Moreno et al. (2015) | From supplementary spreadsheet |

Table 1 – Continued from previous page
| Locality | Strat (m) | Taxon | Source | Comments |
| --- | --- | --- | --- | --- |
| 290466 | NA | NA | Moreno et al. (2015) | From supplementary spreadsheet |
| 390017 | 5 | *Pliomegatherium lelongi* | Amson et al. (2016); Carrillo et al. (2018) | In referred material of Amson et al. without strat position; strat position said to be in the stri database. In table 13 of Carrillo et al. |
| 390018 | 4.5 | Proterotheriidae indet. | Carrillo et al. (2018) | In table 13 |
| 390020 | 5 | Proterotheriidae indet., Toxodontinae indet. | Carrillo et al. (2018) | In table 13 |
| 390022 | NA | Tardigrada indet. | Amson et al. (2016) | In referred material without strat position. Strat position said to be in the stri database |
| 390023 | 3.5 | Lestodontini indet. | Amson et al. (2016); Carrillo et al. (2018) | In Amson et al. in referred material without strat position; strat position said to be in the stri database. In table 13 of Carrillo et al. |
| 390024 | 4.5 | cf. *Nothotherium*, *Pliomegatherium lelongi* | Amson et al. (2016); Carrillo et al. (2018) | In referred material in Amson et al. without strat position; strat position said to be in the stri database. In table 13 of Carrillo et al. |
| 390025 | 4.5 | Scelidotheriinae indet. | Amson et al. (2016); Carrillo et al. (2018) | In Amson et al. in referred material without strat position; strat position said to be in the stri database. In table 13 of Carrillo et al. |
| 390026 | 5 | *Pliomegatherium lelongi* | Amson et al. (2016); Carrillo et al. (2018) | In referred material in Amson et al. without strat position; strat position said to be in the stri database. In table 13 of Carrillo et al. |
| 390075 | 4 | Elasmobranchii | Carrillo-Briceño et al. (2019) | Locality and position previously discussed in Ballen et al. (in prep). Supplementary table S1 of Carrillo-Briceño, no strat position |

Table 1 – Continued from previous page
| Locality | Strat (m) | Taxon | Source | Comments |
| --- | --- | --- | --- | --- |
| 390077 | 1.7 | Elasmobranchii, Megalonychidae sp. nov. | Amson et al. (2016); Carrillo et al. (2018); Carrillo-Briceño et al. (2019) | In referred material of Amson et al. without strat position; said to be in the stri database. In table 13 of Carrillo et al.. In supplementary table S1 without strat position of Carrillo-Briceño et al. |
| 390080 | 5 | Elasmobranchii | Carrillo-Briceño et al. (2019) | Supplementary table S1 of Carrillo-Briceño et al., no strat position |
| 390081 | 21 | NA | Moreno et al. (2015) | From supplementary spreadsheet |
| 390083 | NA | Elasmobranchii | Carrillo-Briceño et al. (2019) | Supplementary table S1, no strat position |
| 430052 | 5 | Proterotheriidae indet., Elasmobranchii | Carrillo et al. (2018); Carrillo-Briceño et al. (2019) | In table 13 of Carrillo et al.. Supplementary table S1 in Carrillo-Briceño et al., no strat position |
| 470059 | 1,7 | NA | Moreno et al. (2015) | From supplementary spreadsheet |
| 470059 | 2 | Elasmobranchii, Toxodontinae indet. | Carrillo et al. (2018); Carrillo-Briceño et al. (2019) | In table 13 of Carrillo et al. Supplementary table S1 of Carrillo-Briceño et al., no strat position |
| 470060 | 4 | Camelidae indet., Lestodontini indet., Megalonychidae sp. nov., Proterotheriidae indet., Toxodontinae indet. | Amson et al. (2016); Carrillo et al. (2018) | In referred material of Amson et al. without strat position; strat position said to be in the stri database. In table 13 in Carrillo et al. |
| 470061 | 4.3 | *Chapalmalania* sp., Proterotheriidae indet., Toxodontinae indet. | Carrillo et al. (2018); Forasiepi et al. (2014) | In table 13 of Carrillo et al. Lower Ware formation in the text, 4.3 in geological setting of Forasiepi et al. |
| 470062 | 4 | NA | Moreno et al. (2015) | From supplementary spreadsheet |

Table 1 – Continued from previous page
| Locality | Strat (m) | Taxon | Source | Comments |
| --- | --- | --- | --- | --- |
| 470062 | 5 | Crocodylidae<br>indet., *Crocodylus* sp., Elasmobranchii,<br>Mylodontidae<br>indet., Proterotheriidae<br>indet. | Amson et al. (2016); Carrillo et al. (2018); Carrillo-Briceño et al. (2019); Moreno-Bernal et al. (2016) | In referred material of Amson et al. without strat position, strat position said to be in the stri database. In table 13 of Carrillo et al. Supp table S1 of Carrillo-Briceño et al. In Moreno-Bernal et al. said to be Police station, no explicit strat position, although the illustrated position in figure 1C points to just below the third conglomerate of the stratotype |
| 470064 | NA | Elasmobranchii | Carrillo-Briceño et al. (2019) | Supplementary table S1, no strat position |
| 600001 | NA | NA | Moreno et al. (2015) | From supplementary spreadsheet |
| NA | ca. 4.8 | *Brachyplatystoma*<br>cf. *vaillantii* | Aguilera et al. (2013) | Said to be from the upper Castillets = Ware. Without strat position data, without STRI locality data |
| NA | ca. 4.8 | Doradidae indet. | Aguilera et al. (2013) | Said to be from the upper Castillets = Ware. Without strat position data, without STRI locality data |

**Figure 1:**
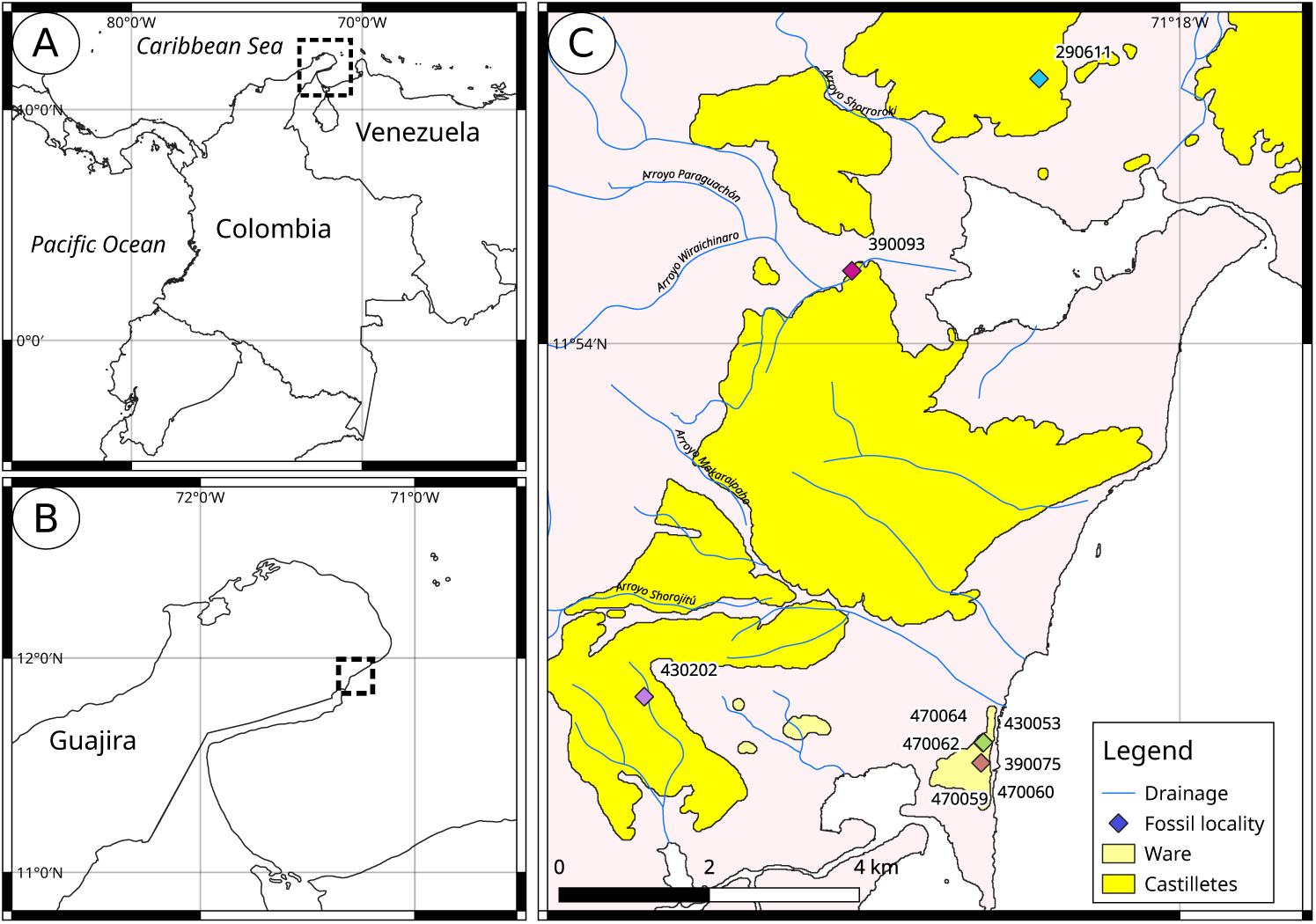
Cartography of the Guajira Peninsula.

**Figure 2:**
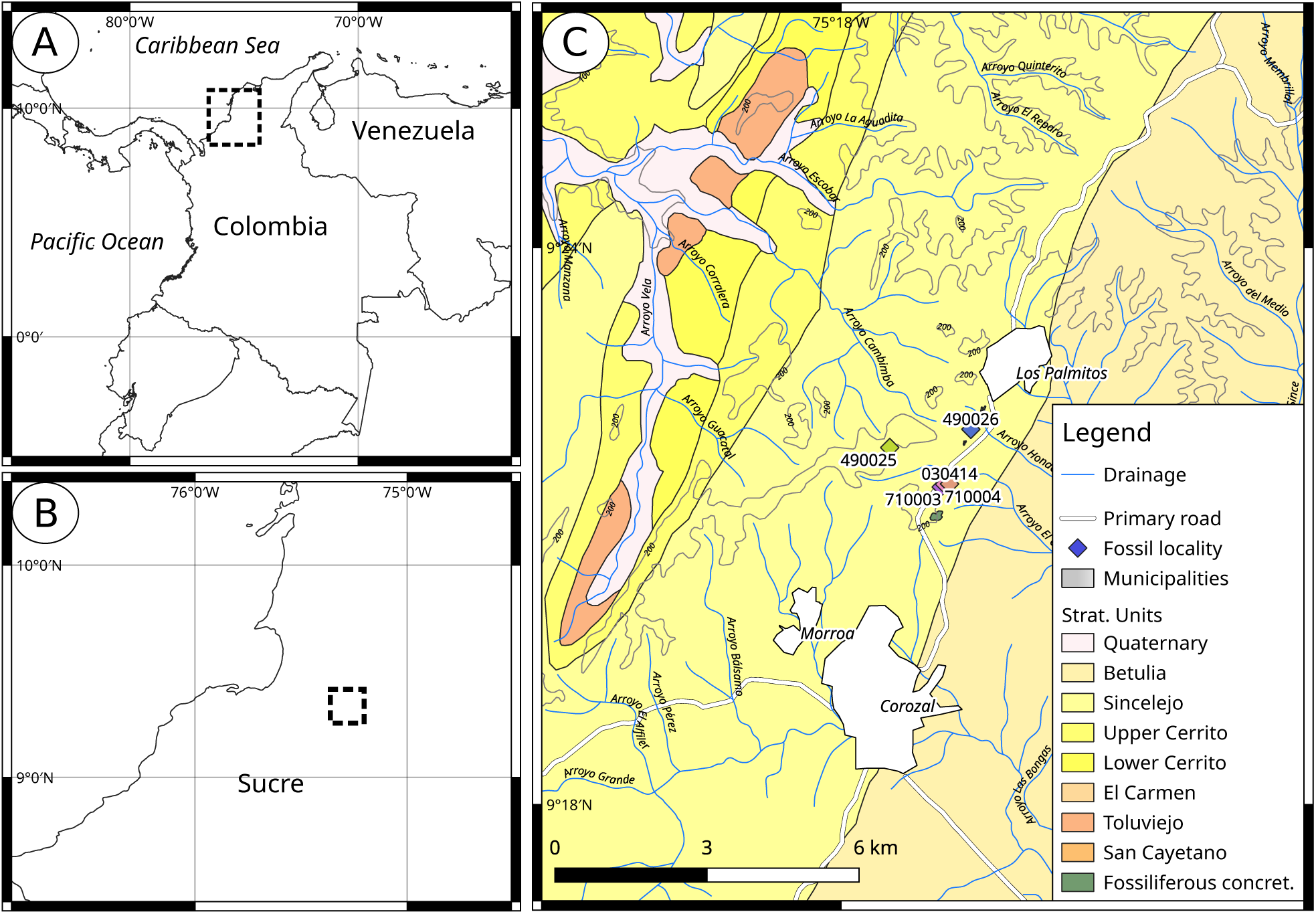
Cartography of the Corozal area.

**Figure 3:**
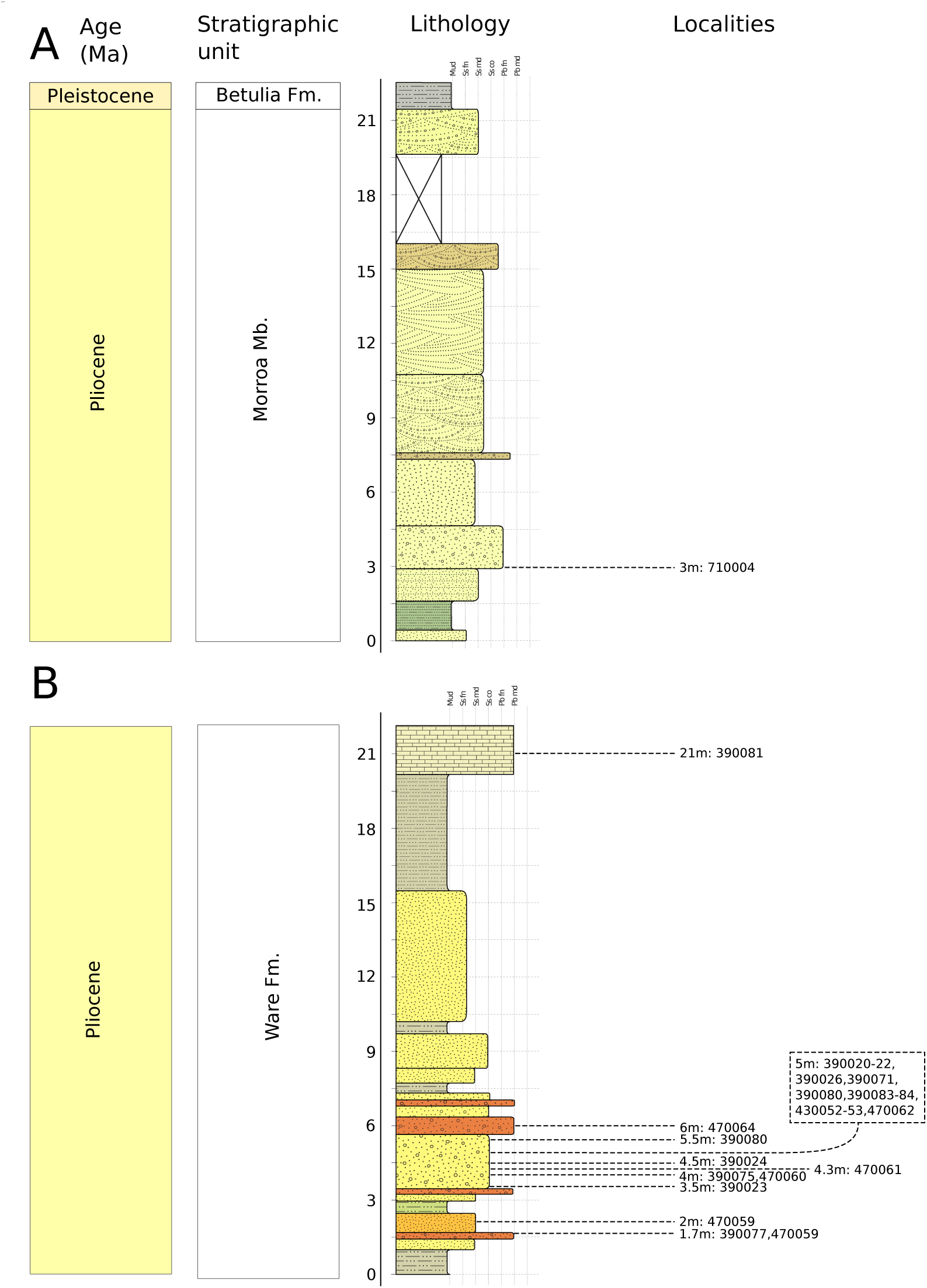
Stratigraphic position of the fossil specimens herein studied. A) San Francisco farm section. B) Ware formation stratotype. Staff scales in stratigraphic meters. Vertical guides are granulometry; Mud = mud, Ss = sand, Pb = pebbles, fn = fine, md = medium, co = coarse. Localites are labeled as with the stratigraphic position in meters from base of the sequence and then the list locality numbers. Colour reflects the one recorded in the outcrop. Stratigraphic columns modified from Moreno et al. (2015)

#### 3.4.2 The San Jacinto tectonic belt

The San Jacinto tectonic belt is an accretionary feature that forms the Cordillera de San Jacinto, a mountain range to the west of the Magdalena-Cauca drainage. It is bound along its east and west margins by the Sinú and Romeral lineaments (Duque-Caro, 1984). This structure and the associated Sinú belt comprise the Sinú–San Jacinto foldbelt that was formed by contraction between the Caribbean Oceanic and the South American crusts, forming north– south-trending folds. It has been subject to a sequence of uplift events during the Cenozoic, with an alternation of unconformities and mostly marine sedimentary units, while the uppermost one is composed of terrestrial sediments. Duque-Caro (1984) calls “Sincelejan Stage” to the association of continental sediments of Pliocene age around Sincelejo between the pre-Late Pliocene and Pleistocene tectonic emergences affecting the San Jacinto tectonic belt. These two events are evidenced by the uncomformities bounding the continental sediments of the Sincelejo formation.

The Sincelejo formation has long been known to preserve fossil vertebrates; however, very few taxa have since been reported in press, and they were restricted to mammals until the present contribution. The first mention of fossil vertebrates from the Sincelejo formation was the report of fossil remains from the Corozal–Sincelejo area by Royo y Gomez (1946, p.499) of reptile bone fragments and a tooth of the Rodent later described as *?Gyriabrus royoi* by Stirton and R. (1947) from the Sierra Peñata locality of the San Antonio(?) sandstone. Stirton (1953) acknowledges the complex stratigraphic of the area, where the names Savana, Sincelejo, and San Antonio have been applied to sandstone facies cropping out in the vicinity of Sincelejo, all with a poor understanding of lateral extent, age, and regional correlation. Stirton however, argues that these sandstone beds might be rather recent based on evolutionary trends in tooth morphology of the Dinomyidae. de Porta (1962), DuqueCaro (1966), and Duque-Caro (1967) focused on the marine facies underlying the Sincelejo formation and studied extensively the molluscs and microfossils, mentioning only succinctly the presence of vertebrates in the continental facies already reported in publications; their descriptions of geographic features make it an important reference for referencing geological features and localities in this area, where the official cartography seems to lack some detail. All these references consistently avoid the nomenclatural problem of the stratigraphy in the region by referring to the sediments above the Cerrito formation with the general term “continental series”. Marshall et al. (1983) mention the fossil record of the Sincelejo area as the Sierra Peñata fauna, and reports the presence of a Toxodont citing de Porta (1961) as the source. The latest review of the fossil assemblages of northern Colombia was carried out by Villarroel and Clavijo (2005), who also described the Glyptodont *Neoglyptatelus sincelejanus* from a locality 6 km south of Sincejelo in the Calle Fría–Segovia road. These authors also reported indeterminate remains of a Toxodont from 2 km to the north of Los Palmitos, in the vicinity of Corozal, and the Toxodont cf. *Trigodonops* from sandstone beds where the town of Corozal was built. These authors recognize with some uncertainty the Sincelejo formation as the source of the indeterminate Toxodont and cf. *Trigodonops* mentioned above, and an uncertain unit below the Sincelejo formation as the source of *Neoglyptatelus sincelejanus* and *?Gyriabrus royoi* (see figure 3, p.354), also challenging the local stratigraphy of previous works on the basis of field observations. Villarroel and Clavijo (2005) however, did not cite Clavijo-Torres and Barrera-Olmo (2001) and therefore can not be correlated directly to the latter work that has been followed by later authors in recognizing the Betulia, Sincelejo, and Cerrito as distinct formations with more or less clear stratigraphic boundaries.

#### 3.4.3 Fossil localities

Most of the vertebrate localities from the Ware formation are very spatially restricted, occurring in a small area in the stratotype of the unit. The main difference between localities is the fine-scale stratigraphic provenance, sometimes to fractions of stratigraphic meters. Several localities have been named differently when in fact they are synonyms as the main criterion for locality designation in the field was stratigraphic position. Most of the explicit, textual information about stratigraphic position of the different STRI localities come from Amson et al. (2016) and Carrillo et al. (2018), while other references have either cited any of these sources, the original paper by Moreno et al. (2015), or stated the STRI sample online database as online repository of provenance information.

A compendium of published localities in the stratotype of the Ware formation is provided in Table 1, along with a synonymy of the stratigraphic section (Figure 3). As the original catalog information sometimes suffers from synonymous locality labeling and the ultimate criterion for locality naming is stratigraphic position, some of the original localities from the same fossil level bear two locality names. I retain the original locality labels as they are expected to remain attached to the associated provenance data and field notes from different collectors (e.g., Figure 16). The present compendium of locality information is expected to serve as permanent future reference.

## 4 Results

### 4.1 Systematic Palaeontology

> **Teleostei Ostariophysi Characiformes Anostomidae**
>
> **Genera Leporinus or Hypomasticus**
>
> **Gen. et. sp. indet.**
>
> Figure 4.

**Figure 4:**
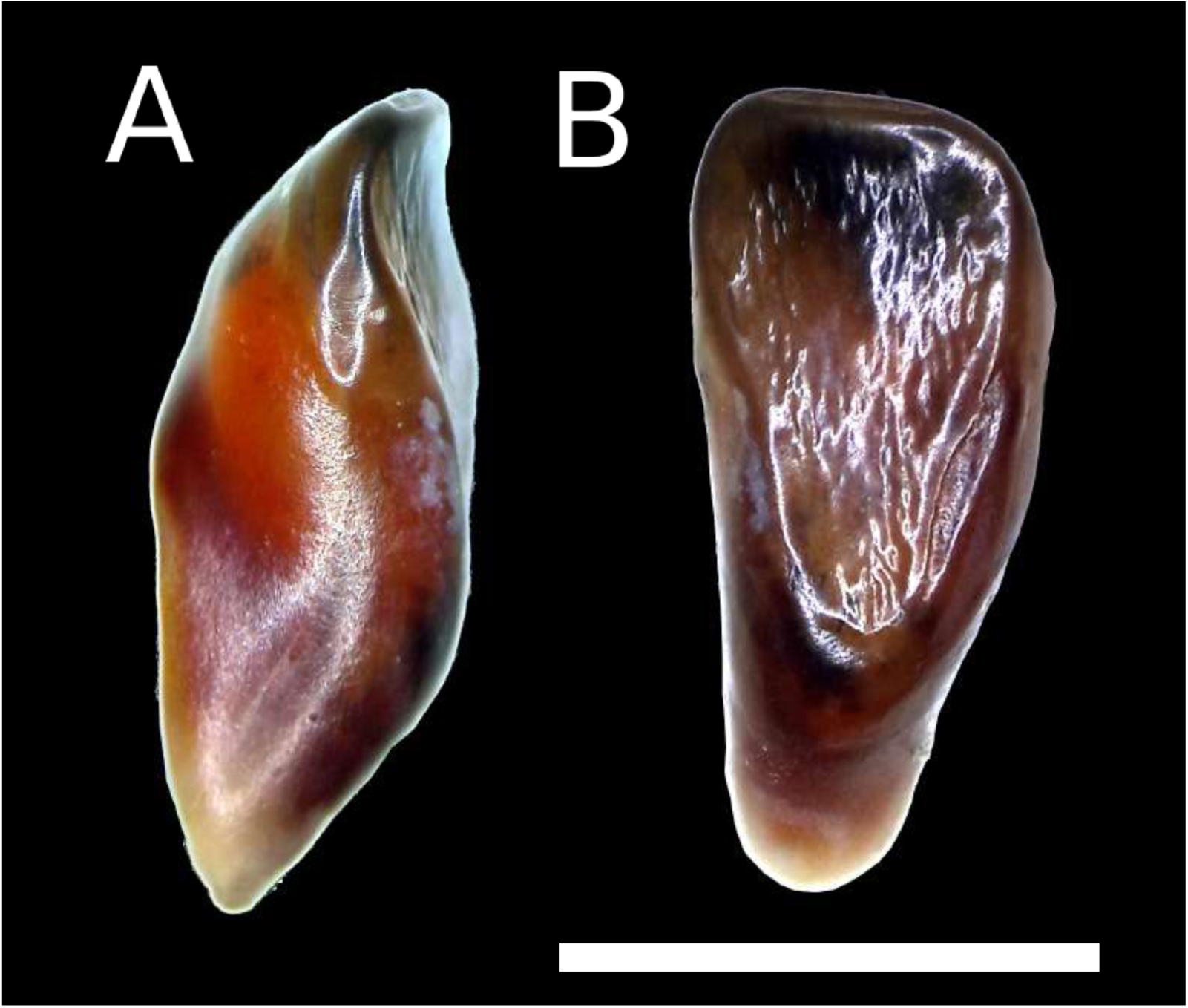
Anostomid tooth from the Sincelejo formation. A) Lateral view. B) Lingual view. Scale bar equals 5 mm.

**Provenance**—Sincelejo formation, Locality STRI 710004, GAB-P 415, isolated symphysial dentary tooth.

**Description**—Isolated tooth lacking strong asymmetry, unicuspid, without strong distal projection. Crown elongate, not wider than long. Occlusal surface striate somewhat inverted triangular in outline.

**Remarks**—The isolated tooth can be assigned with confidence to family and has been further identified as a member of either *Leporinus* or *Hypomasticus* (as defined by Sidlauskas and Vari, 2008). Those genera consist of strictly cis-Andean (i.e., east of the Andes) species, so this record further reinforces a relationship between the Magdalena-Cauca drainage with cis-Andean drainages by the Pliocene. The genus *Megaleporinus* was recently separated from *Leporinus* to accommodate ten species formerly considered as either *Leporinus* (9 spp.) or *Hypomasticus* (1 sp.) (Ramirez et al., 2017). Species of *Megaleporinus* are mostly cis-Andean with one member found in trans-Andean (i.e., west of the Andes) drainages (*Megaleporinus muyscorum*), and have dentary teeth that are very wide, semispherical, and spoon-shaped (e.g., fig. 2, Ramirez et al., 2017) in contrast to the fossil specimen which is clearly elongate and not spoon-shaped. *Hypomasticus* is a problematic genus whose delimitation is still confusing, and *Leporinus* is a large genus whose alpha-taxonomy has improved considerably in the last few decades. Further study of recent material of those genera as well as a better understanding of their respective delimitation are needed to evaluate the relationships of the specimen.

> **Family Serrasalmidae**
>
> **Genus *Serrasalmus* Lacépède, 1803**
>
> **Serrasalmus sp.**
>
> Figure 7.

**Figure 5:**
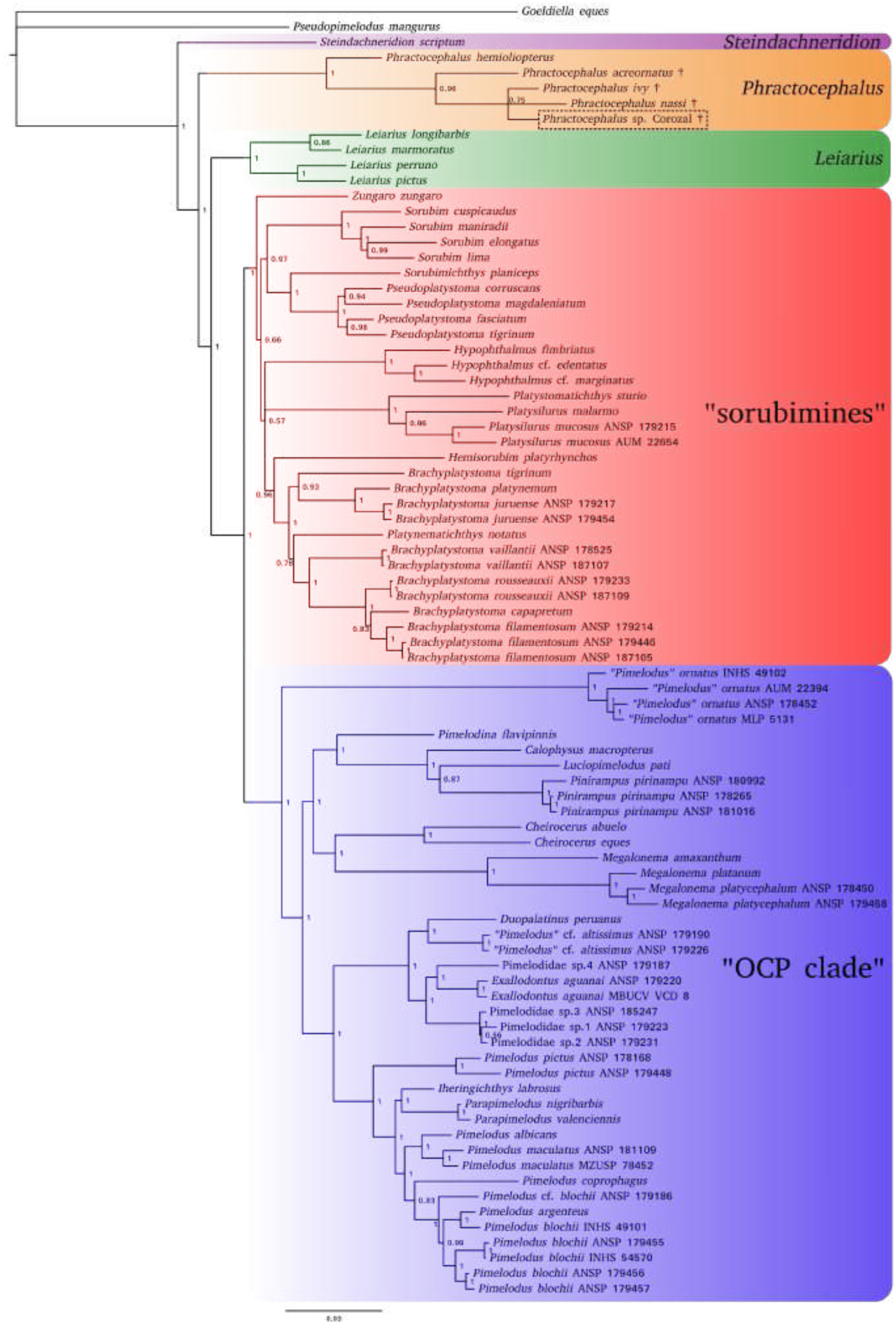
Bayesian phylogenetic analysis of the Pimelodidae including fossil representatives of the genus *Phractocephalus*. Nodal support values are bayesian posterior probabilities. Dagger (†) indicates extinct lineages while the fossil occurrence from the Sincelejo Formation is enclosed in the dashed box. Relevant monophyletic groups are highlighted in colours.

**Figure 6:**
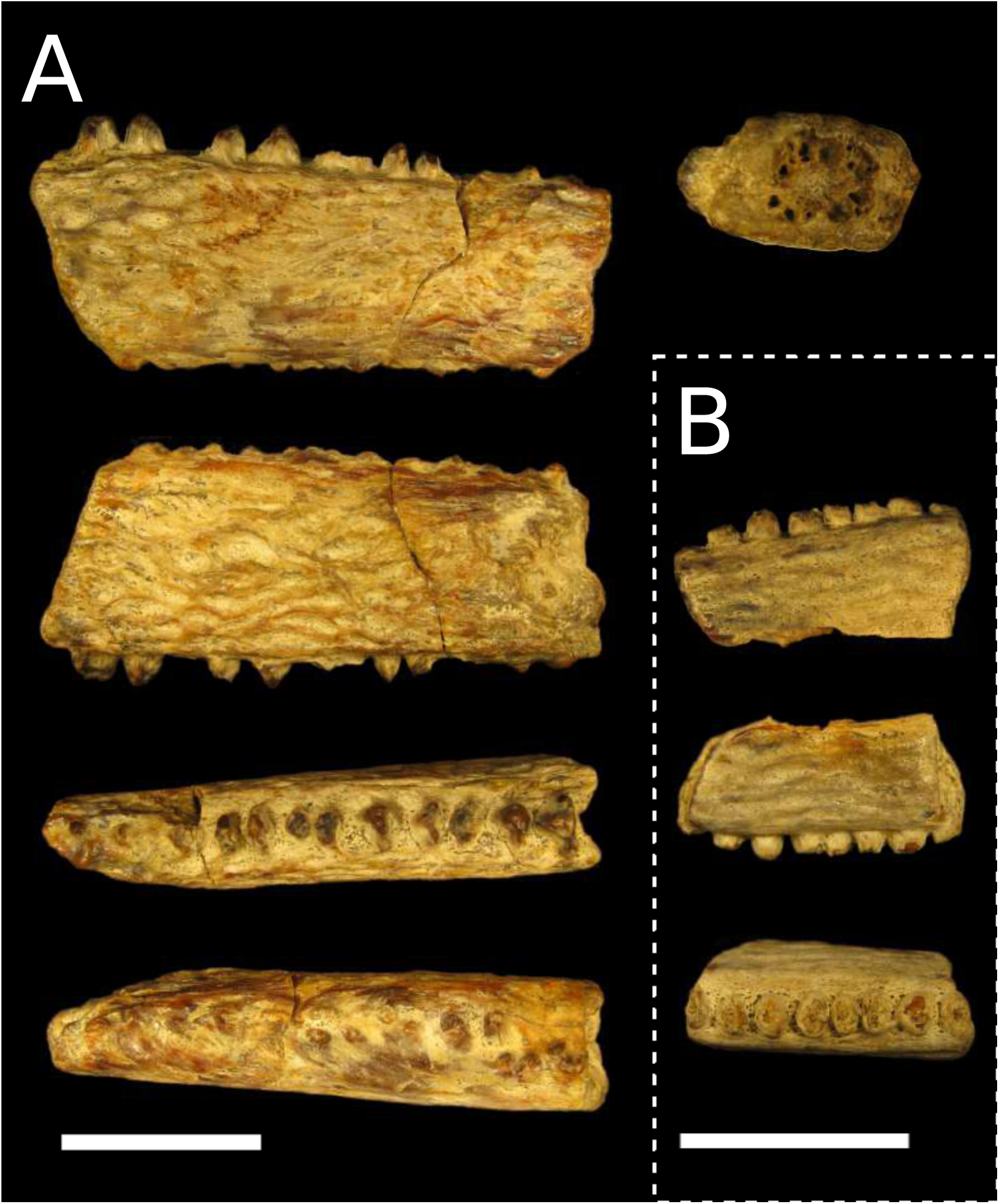
Pectoral-fin spines of *Phractocephalus* sp. A) Fossil pectoral spine fragment, MUN 41058, specimen in dorsal, ventral, anterior, posterior, and proximal cross-section views. B) Fossil spine fragment, MUN 43679, specimen in dorsal, ventral, and anterior views. Scale bar equals 20 mm in all cases.

**Figure 7:**
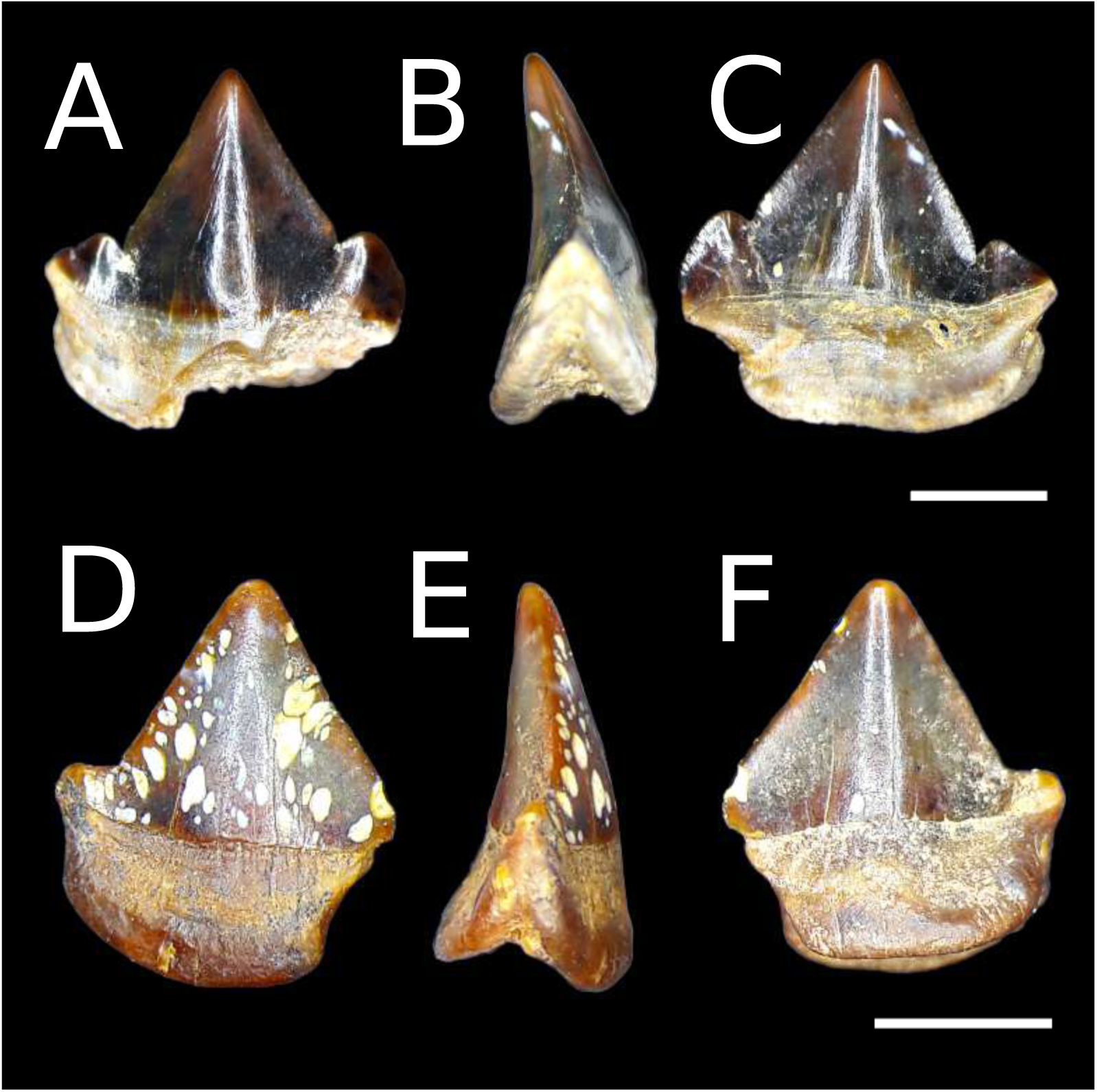
Fossil isolated teeth of *Serrasalmus* sp. A-C) Right D1 tooth in labial, symphysial, and lingual views respectively. D-F) Left D4-D5 tooth in labial, commisural, and lingual views respectively. Scale bars equal 2 mm.

**Provenance**—Ware formation, Locality STRI 470060, MUN 34443, one isolated symphysial dentary (D1) tooth. Locality STRI 470059, MUN 37712, one dentary tooth D3-D4, almost complete.

**Description:** Left symphysial dentary tooth labio-lingually compressed, crown slightly curved lingually, tricuspid with main cusp largest and lateral cusps about same size, cutting edge very sharp. Root bifid in commisuro-symphysial view, root halves in angle of ca. 40*^◦^*, basal outline smoothly straight to slightly concave in middle section. Right dentary tooth D3-D4 labio-lingually compressed, crown almost completely triangular occlusal outline in labial view and slightly curved lingually, bicuspid, with main cusp largest and very small commisural cusp somewhat deflected comisurally. Root bifid in commisuro-symphysial view, root halves at angle of ca. 55*^◦^*, basal outline smoothly convex in labial view.

**Remarks**—It is possible to distinguish between premaxillary teeth from dentary teeth in carnivore serrasalmids given that the premaxillary ones have either a lingual projection that forms an occlusal platform similar to the molariform platform in herbivorous pacus or are much more asymmetric than dentary teeth; oftentimes, the premaxillary teeth have very wide base and low crown, with the commisural cusplet more developed than in dentary teeth. Also, premaxillary teeth have a less-developed inter-dental anchoring system when compared to dentary teeth where such anchoring mechanism is less developed.

Dentary symphysial teeth in carnivore piranhas (i.e., *Pygocentrus* and *Serrasalmus*) are tri- to penta-cuspidate and lack the commisural socket present in subsequent dentary teeth for anchoring of the anterior adjacent tooth. In *Pygocentrus* the cusplets around the main one are strongly heterogeneous in size, being the commisural cusplet larger than the symphysial one; on the contrary, these cusplets are about the same size in *Serrasalmus*. Other Serrasalmids with multicuspidate teeth present either mamilliform (*Catoprion*) or multicuspidate with four or more cusplets (*Pristobrycon* and *Pygopristis*).

The identification of the specimens herein studied as dentary teeth of the genus *Serrasalmus* is based on the lack the occlusal platform seen in premaxillary teeth, the presence of cusplets on both sides of the main cusplet which is found in symphysial teeth, and the the presence of cusplet of about the same size that allows to identify the genus *Serrasalmus*.

The tricuspidate condition in piranha teeth is also seen in the fifth premaxillary tooth of the genus *Pygocentrus* (*vs.* fifth premaxillary tooth bicuspid in *Serrasalmus*). As already mentioned the symphysial teeth of either the premaxilla or the dentary can also be tricuspid, have narrow base, and have a nearly symmetrical main cusplet; in contrast, the fifth premaxillary tooth in *Pygocentrus* shows a very wide base and asymmetrical main cusplet which is commisurally-oriented. Lundberg (1997) illustrated this morphology from an isolated tooth IGM 251277 from the middle Miocene La Vent fauna in central Colombia; although the author was uncertain about the affinities of such specimen and thus identified it as “*Serrasalmus*, *Pygocentrus*, or *Pristobrycon* sp.”, it is possible to narrow down its identity as a fifth premaxillary tooth of the genus *Pygocentrus* sp.

Antoine et al. (2016) described an extensive collection of vertebrate fossil remains from Contamana in Peru. Among these specimens is a serrasalmid symphysial tooth illustrated in their figure 8A. This specimen can be identified as a representative of the *Pygocentrus* using the diagnostic characters discussed above. This specimen represents the oldest exemplar of the genus in the fossil record, as it was collected at the locality CTA-32, above the unconformity to the base of the Chambira formation and below the dated level CTA-08SA (26.56 ± 0.07 Ma).

**Figure 8:**
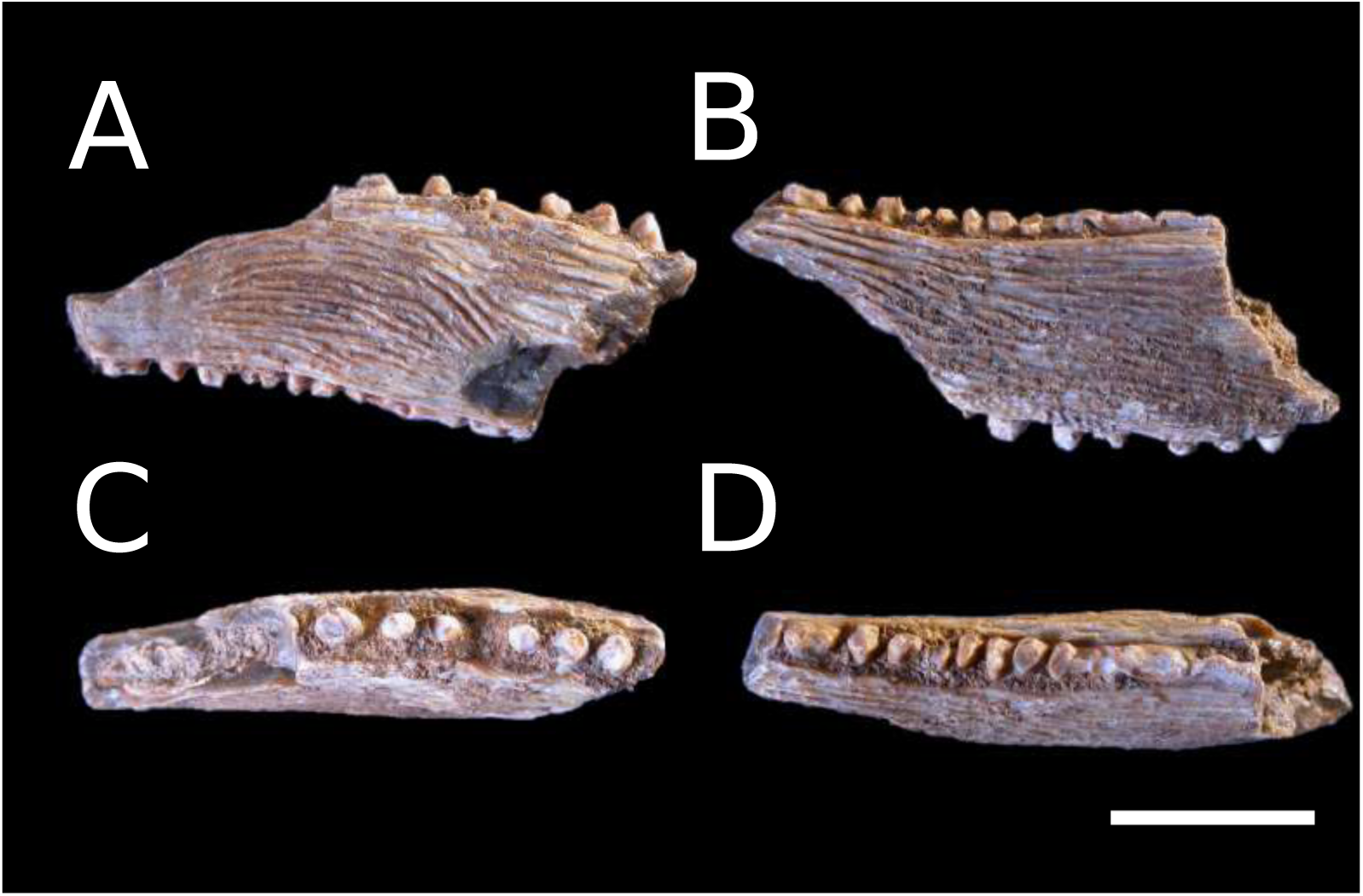
Left pectoral-fin spine of *Trachelyopterichthys* sp., MUN 34401. A-D Dorsal, ventral, anterior, and posterior views respectively. Scale bar equals 10 mm.

> **Order Siluriformes Family Auchenipteridae**
>
> **Genus Trachelyopterichthys Bleeker, 1862**
>
> **Trachelyopterichthys sp.**
>
> Figure 8.

**Provenance**—Ware formation, Locality 470060, MUN 34401, one pectoral-spine fragment.

**Description**—Left pectoral-spine fragment preserving the middle portion of the spine shaft along with the anterior, posterior, dorsal, and ventral ornaments. Shaft depressed and robust, with antero-dorsal, antero-ventral, postero-dorsal, and postero-ventral margins smooth. Preserved shaft showing some curvature along the axis posteriorly. Dorsal and ventral ornament consisting of undulating, subparallel ridges, decreasing in thickness from anterior to posterior across shaft. Abundant tubercles onto the dorsal ridges throughout the shaft, ventral tubercles present, yet restricted to the anterior 1/4 of ventral surface. Anterior ornament consisting of straight to slightly antrorse spinules directly implanted onto anterior shaft surface. Posterior ornament consisting of irregular blades poorly preserved but showing extensive fusion between units, straight to slightly retrorse in orientation. Lumen comprising ca. 1/2 of shaft area in cross-section; lumen outline very irregular.

**Remarks**—The anterior ornament of the pectoral-fin spine in *Trachycorystes* consists of straight spinules with fused bases, so that the whole ornament series consists of a continuous ridge emerging from the anterior surface in dorsal view. In contrast, the anterior spinules are sessile and implanted directly on the anterior shaft surface without fusion at the bases in *Trachelyopterichthys*. This is the first fossil record of the family Auchenipteridae anywhere. The family has a Neotropical distribution in drainages on both sides of the Andes, with a greater diversity in cis-Andean basins. The genus *Trachelyopterichthys* is composed of only two species: *T. anduzei* from the Orinoco drainage, and *T. taeniatus* from the Amazon drainage (Birindelli, 2014; Calegari et al., 2019). Further differences between species are not known, but further study of extant specimens of the genus may provide further information on the identity of the fossil as belonging to either of the extant species or even an extinct, undescribed one. At present it was not possible to reach at a conclusion as to the specific identity of the fossil remains.

> **Family Doradidae**
>
> **Genus *Hemidoras* Bleeker, 1858**
>
> ***Hemidoras* sp.**
>
> Figure 9.

**Figure 9:**
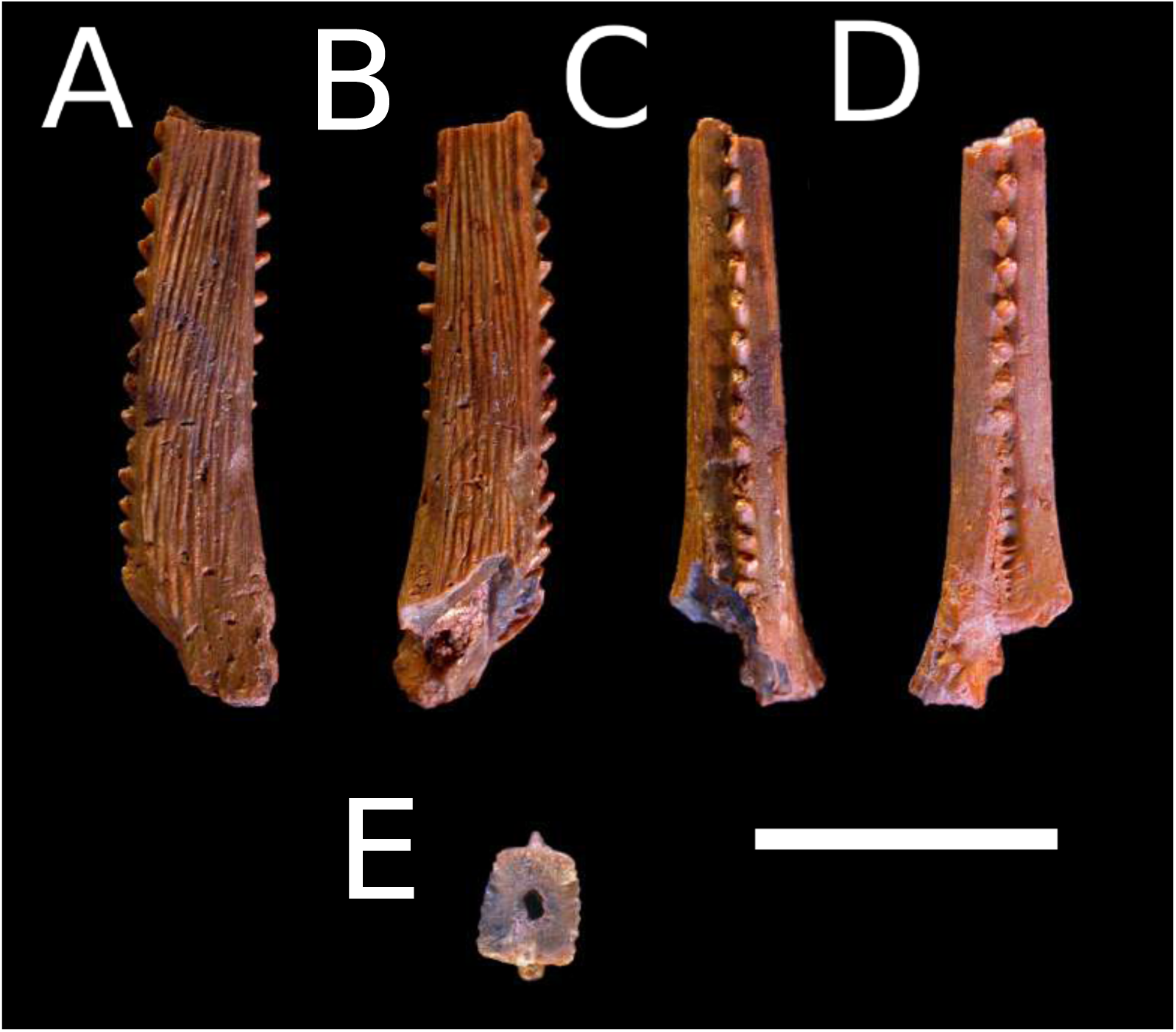
Dorsal-fin spine fragment of *Hemidoras* sp., MUN 34455. A-D) Left, right, anterior, and posterior views respectively. E) Shaft cross section. Scale bar equals 10 mm.

**Provenance**—Ware formation, Locality 470060, MUN 34455, one partial dorsal-fin spine preserving the a left portion of the base and about 1/3–1/2 of the shaft.

**Description**—Base preserving half inflection point of anterior longitudinal ridge, and left lateral condyle but without lateral articular surface; posterior process not preserved. Shaft preserving anterior, posterior, and lateral ornaments, as well as fenestrae communicating interior of shaft lumen with epidermal tissue. Anterior ornament consisting of antrorse spinules from inflection point of anterior longitudinal ridge up to preserved dorsalmost portion of shaft; spinules closely set, leaving space inbetween less than spinule diameter. Posterior ornament absent on basal third of preserved spine length, then consisting of antrorse spinules with space inbetween measuring equaling spinule diameter or more. Lateral ornament consisting of subparallel ridges oriented somewhat oblique to shaft main axis; ridges with angular surface and flat sides. Shaft outline quadrangular in transversal view; lumen maximum axis comprising 1/4 of shaft width in anteroposterior axis.

**Remarks**—The presence of both anterior and posterior antrorse dorsal-fin spine ornaments is a rare feature in the Doradidae, found only in the genera *Hemidoras*, one species of *Leptodoras* (*L. acipenserinus*), one species of *Nemadoras* (*N. humeralis*), *Ossancora*, *Oxydoras*, *Petalodoras*, one species of *Platydoras* (*P. armatulus*), and *Trachydoras*. In contrast, most other taxa show anterior antrorse and posterior retrorse or straight ornaments. Among those taxa, posterior ornaments are separated by a large space of more than 1.5 of the adjacent ornament base width in *L. acipenserinus*, *Ossancora*, *Oxydoras*, *Petalodoras*, *Platydoras*, and *Tenellus*. *Nemadoras humeralis* shows very closely-set posterior ornaments separated by a space of ca. 1.2 of ornament base width. This is in contrast to *Hemidoras* and the fossil specimen where it is between 1.0 and 1.5. Species of *Trachydoras* have extensive posterior ornament developed down to the level of the dorsal-fin spine base, while in the fossil and *Hemidoras* there is at least an extension of 1/4 of the spine shaft length devoid of ornaments basally. This combination of characters allows positive identification of dorsal-fin spines to *Hemidoras* among doradids.

Birindelli (2014) considered *Opsodoras* as a synonym of *Hemidoras*, comprising now five species distributed through the Amazon region in the Amazonas main channel as well as in the basins Beni-Madre de Dios, Branco, Guaporé, Japurá, Middle-lower Madeira, Mamoré, Marañon-Nanay, Negro, Purus, Putumayo, Tefé, Trombetas (Dagosta and de Pinna, 2019), all in cis-Andean South America. Phylogenetic analyses consistently recover *Hemidoras* as part of a group including *Anduzedoras*, *Doras*, *Ossancora*, *Nemadoras*, *Hassar*, *Trachydoras*, and *Leptodoras*; however, the interrelationships among these genera sometimes referred to as the “*Doras* clade” or the “fimbriate-barbel doradids” are variable depending on both the data source (either morphology or sequence data) and the analytical strategy employed (ML, bayesian inference, or parsimony). Arce H et al. (2013, fig. 2) considered *Hemidoras* monophyletic only with the inclusion of *Opsodoras*. The two genera were later synonymized by Birindelli (2014, fig. 70). Birindelli also recovered *Hemidoras* as monophyletic (after inclusion of *Opsodoras*) and sister to a clade comprising *Nemadoras*, *Tenellus*, *Hassar*, *Anduzedoras*, and *Leptodoras*. This conclusion was reiterated by Sabaj and Arce H. (2021).

> **Genus *Oxydoras* Kner, 1855**
>
> **cf. *Oxydoras* sp.**
>
> Figure 10.

**Figure 10:**
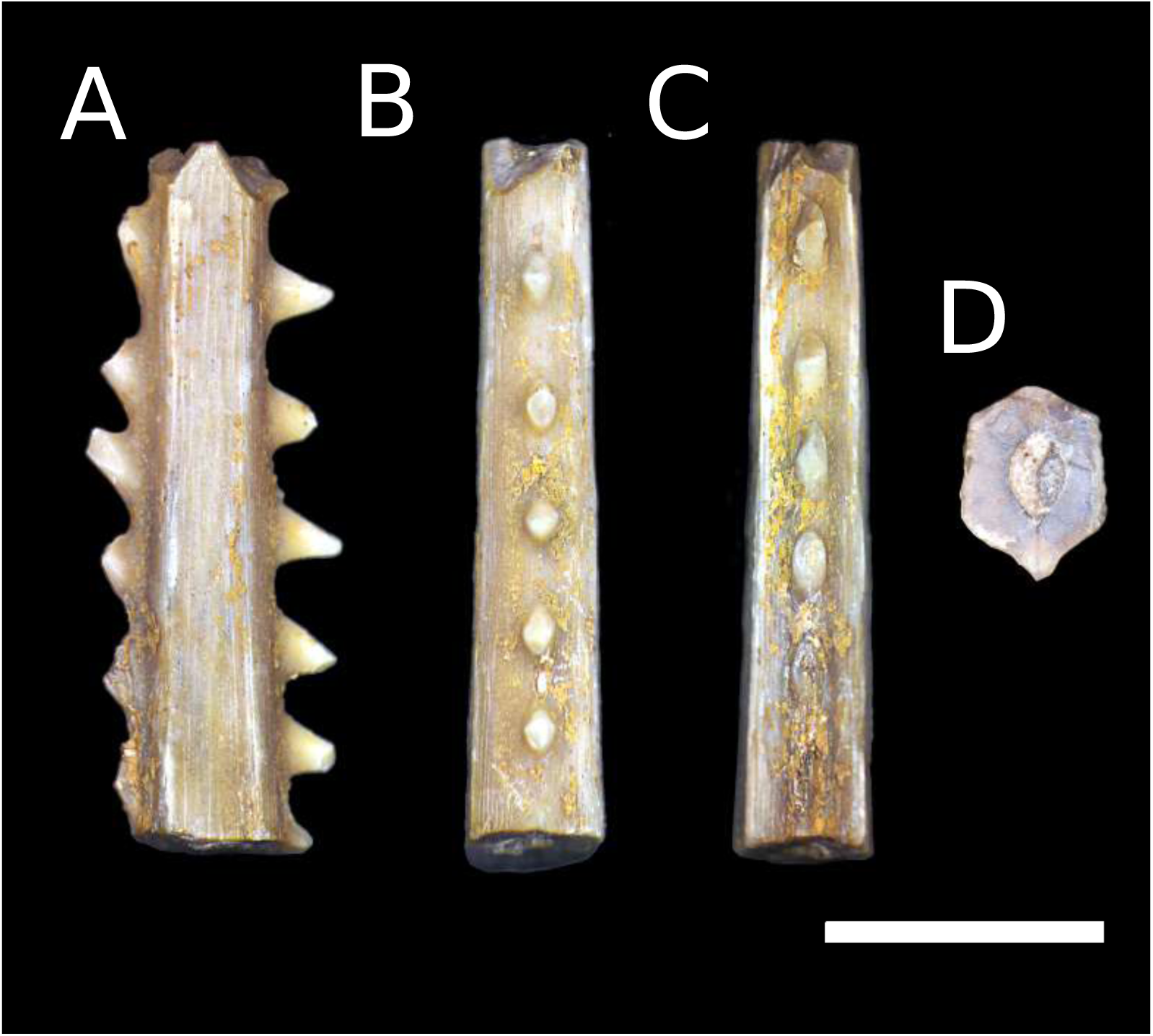
Dorsal-fin spine fragment of cf. *Oxydoras* sp., MUN 34409-2. A-C) Left, posterior, and anterior views respectively. D) Shaft cross section. Scale bar equals 5 mm.

**Provenance**—Ware formation, Locality 470060, MUN 34409-2, distal portion of a dorsal-fin spine shaft.

**Description**—Shaft preserving anterior, posterior, and lateral ornaments. Anterior ornament consisting of antrorse spinules along preserved shaft surface; spinules irregularly spaced, leaving space inbetween less than spinule diameter, fused basally in a longitudinal ridge. Posterior ornament consisting of straight to retrorse spinules with large space inbetween larger than 1.0–1.5 times spinule diameter. Lateral ornament almost absent, with very smooth surface and just faint indication of longitudinal small ridges. Shaft outline hexagonal in transversal view; lumen maximum axis comprising 1/2 of shaft width in anteroposterior axis.

**Remarks**—The genus *Oxydoras* is peculiar among doradids in having a very slender dorsal-fin spine with very spaced out anterior and posterior ornaments. Species of *Oxydoras* have almost smooth lateral shaft surface lacking prominent ridges, unlike all the remaining genera in the family that show lateral shaft surfaces with ridges of varying degrees of development, from fine and numerous as in *Hemidoras* to few in number but hypertrophied as in *Petalodoras* and members of the Astrodoradinae. The fossil specimen herein described matches representatives of the genus *Oxydoras* in most details, except for the orientation of the posterior dorsal-fin spine ornament which is antrorse to straight in most species of *Oxydoras* and retrorse in the fossil specimen. *Oxydoras sifontesi* differs further from the fossil specimen due to the presence of small tubercles on the lateral shaft ornament restricted to the anteriormost third on the spine, while such tubercles are absent in other representatives and the fossil specimen. Due to the latter incongruent character the fossil cannot be allocated with certainty to the genus *Oxydoras*.

> **Family Pimelodidae**
>
> **Genus *Brachyplatystoma* Bleeker, 1862**
>
> **Brachyplatystoma cf. vaillantii**
>
> Figure 11.

**Figure 11:**
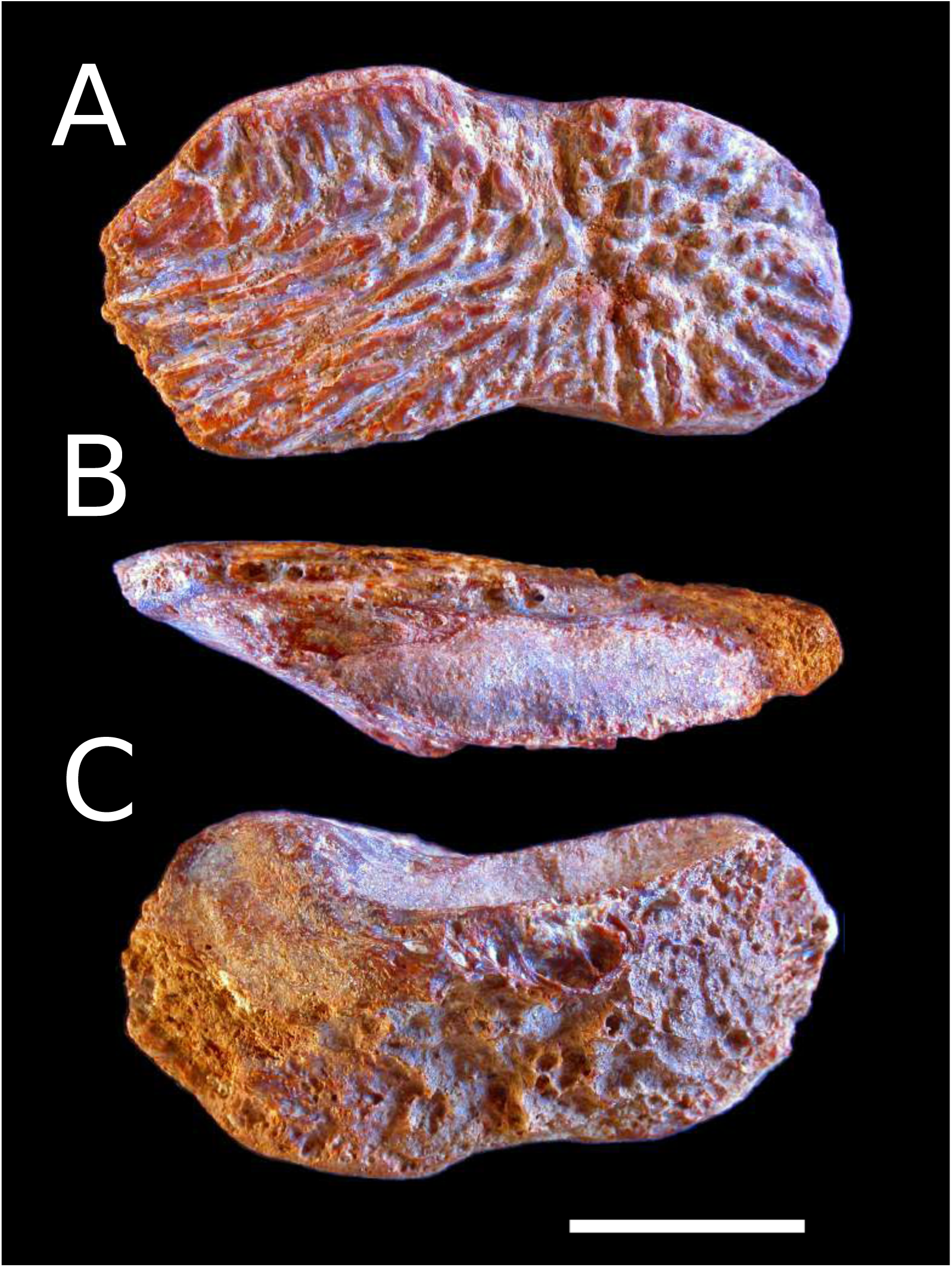
Left sphenotic of *Brachyplatystoma* cf. *vaillantii*, MUN 37567. A-C) Dorsal, lateral, and ventral views respectively. Scale bar equals 10 mm.

**Provenance**—Ware formation, Locality 470059, MUN 37567, partial left sphenotic.

**Description**—Preserving most of the original bone, including the articular facet for the hyomandibula and the surface ornament. Preserved outline renoid in dorsal view. Most of the ventral surface eroded and exposing the internal trabecular structure. Dorsal ornament consisting of parallel ridges on both the lateral surface and the anterior half of the bone in a radiating pattern with focus in the posterior half. Tubercles concentrated in the region of the ridge focus, somewhat aligned to the ridges. Articular facet for the hyomandibula smooth and concave, becoming shallower towards the sides of the bone and suggesting that the posterior portion on the pterotic should have been reduced as most of the functional surface is restricted to the sphenotic.

**Remarks**—*Brachyplatystoma* is a genus with seven extant and one extinct species (Lundberg, 2005; Lundberg and Akama, 2005). Fossil occurrences of this group are known from several localities of Neogene age in northern South America (Lundberg et al., 2010; Aguilera et al., 2013), mostly identified as *Brachyplatystoma* cf. *vaillantii* and *B. promagdalena*. The former species is distinctive among congeners by the pattern of ornamentation on exposed cranial bones, with ridges and tubercles aligned in radiating patterns with focus in the sphenotics. In contrast, other species of the genus have very smooth cranial bones. The only specimen known of *B. promagdalena* is a weberian apparatus, and consequently we lack further anatomical details to compare with congeners. By the distribution of character states, *B. promagdalena* should also show smooth cranial bones, quadrangular opercle, and a well-defined sulcus on the anterior surface of the pectoral spine, the latter two characters being morphological synapomorphies for the genus (Lundberg and Akama, 2005). The numerous Colombian and Venezuelan Neogene occurrences historically associated to *B. vaillantii* must be restudied in order to better assess their specific identity, as they appear to be a different, extinct species sharing the pattern of cranial ornament with the extant *B. vaillantii* (J.G. Lundberg, *pers. comm.*).

> **Genus *Phractocephalus* Spix & Agassiz, 1829**
>
> **Phractocephalus sp.**
>
> Figure 12,6.

**Figure 12:**
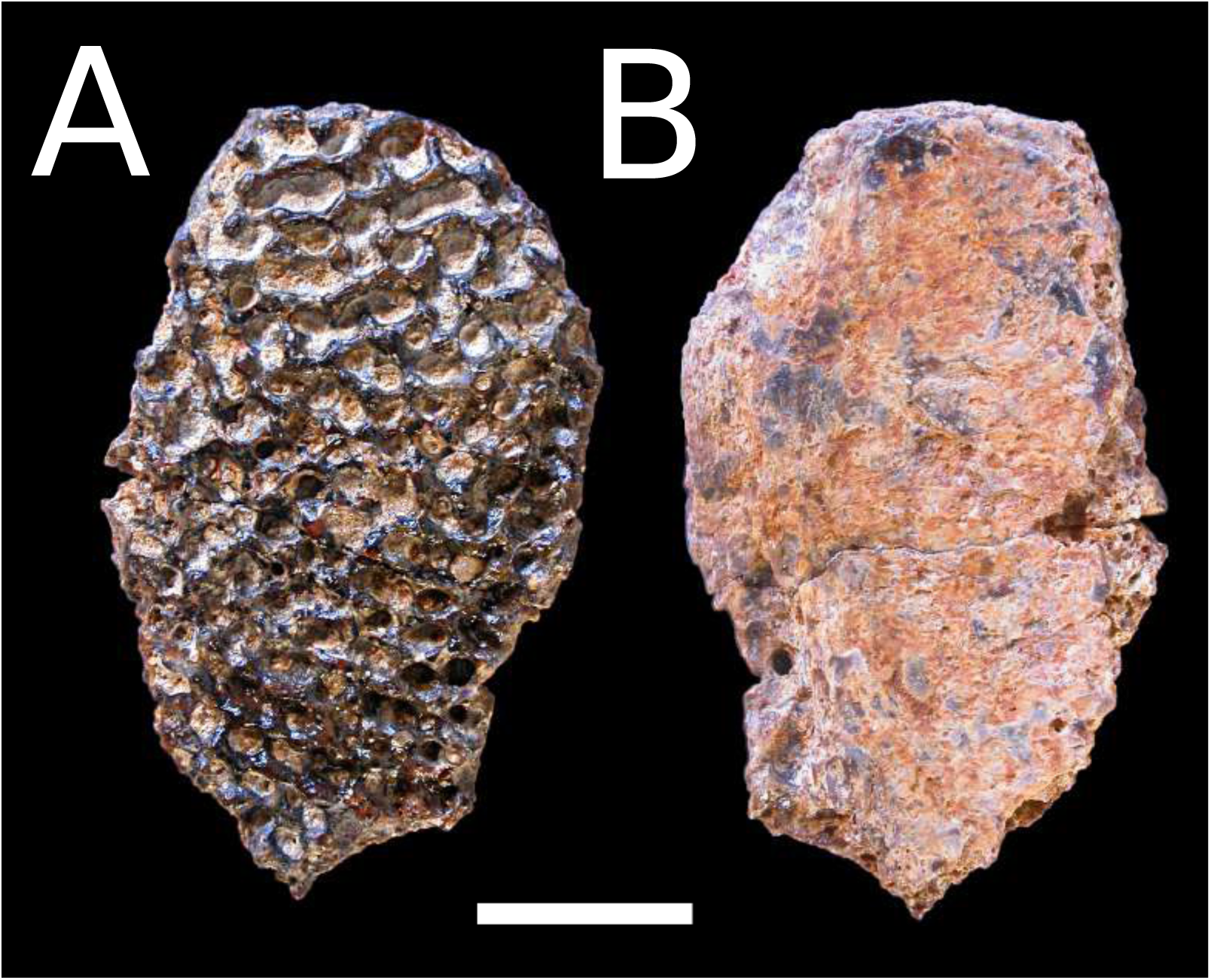
Nuchal plate fragment of *Phractocephalus* sp., MUN 34425. A-B Dorsal and ventral views respectively. Scale bar equals 10 mm.

**Provenance**—Sincelejo formation, Locality STRI 710004, MUN 41058, MUN 43679, pectoral spine fragments. Ware formation, Locality 470060, MUN 34425, partial nuchal plate.

**Description**—The Pectoral-spine fragments preserve the proximal portion of the right pectoral-spine shaft with dorsal, ventral, anterior and posterior ornament preserved, with proximal exposition of the medullar region. Dorsal and ventral ornament consisting of reticulate ridges with ovoid pits; dorsal and ventral ornament appear to be equally well developed. Anterior ornament consisting of straight, vertically-elongate ornament with some tips worn. Posterior ornament consisting of retrorse, oblique spinules directed somewhat ventrally; the ornament disposition is typical of that of ornaments on the basal portion of the spine where the fin membrane tends to insert dorsal to the ornaments, making them to direct slightly to the ventral portion of the spine shaft. Even when the posterior ornament is retrorse in curvature, they tend to be displaced forward, a condition typical of basal ornaments in catfishes of the Pimelodidae. Medullar region in cross-section spongy with a small and oval lumen. MUN 43679 preserves only the anterior portion of the middle region of the right pectoral-spine shaft with dorsal and ventral reticulate ridges; anterodorsal and anteroventral margins smooth; anterior ornament consisting of straight, vertically-elongate ornament, with tips eroded due to transport.

The fragment of nuchal plate (MUN 34425) is oval in outline. No margins preserved, only dorsal and ventral surfaces. Dorsal ornament consisting of strong reticulating ridges surrounding round to ovoid pits. Ventral surface smooth with traces of abrasion and some exposure of spongy bone internal structure.

**Remarks**—The phylogenetic analysis of combined molecular and morphological data for the family Pimelodidae indicates that the specimens of *Phractocephalus* sp. herein described lie in a clade along with *P. nassi* and *P. ivy*, the two extinct species of *Phractocephalus* (0.75), while *P. acreornatus* is recovered as basal to that trichotomy (0.96); the extant species *P. hemioliopterus* is to the most basal member of the genus (Figure 5). Two large traditional groups recognized informally for the family were recovered as monophyletic in our analysis (the so-called ”sorubimines” and the ”OCP clade” of Lundberg et al., 2011), while *Steindachneridion*, *Phractocephalus*, and *Leiarius* were found to be successive groups to the base of the Pimelodidae. Overall the phylogenetic analysis shows high nodal support values with posterior probabilities above 0.95 in almost all cases; nine out of 80 nodes showed posterior probabilities between 0.5 and 0.9. These results place the specimens form the Sincelejo within *Phractocephalus* with high confidence and corroborate earlier hypotheses of the interrelationships of Pimelodid Catfishes (Lundberg et al., 2011; Silva et al., 2024).

The pectoral spines of *Phractocephalus* differ from all other neotropical genera by the presence of reticulated dorsal and ventral surface ornament (vs. spines lacking ornament or with ornament present but never reticulated). Also, pectoral spines of *Phractocephalus* differ from pectoral spines in other pimelodid genera by the presence of vertically-bifid anterior ornament (vs. anterior ornament consisting of vertically-unicuspid ornament or absent), and from other pimelodids except *Leiarius* by the presence of unicuspid and smooth spinules (vs. anterior ornament consisting of tubercles or absent).

The characteristic surface ornament of the nuchal plate consisting of strong reticulating ridges around round to oval pits is a feature that easily distinguishes the genus *Phractocephalus* from other Neotropical Siluriforms; this feature has already been used in the literature when assessing the generic position of bone elements in this genus in Neotropical fossil fish faunas (Lundberg and Aguilera, 2003; Aguilera et al., 2008; Azpelicueta and Cione, 2016). Although catfishes of the families Auchenipteridae, Ariidae, and Doradidae display very developed cranial ornamentation through exostosis, those cases often consist of either subparallel ridges, series of tubercles, or a combination of both, but never arranged in a reticulate pattern.

> **Genus *Platysilurus* Haseman, 1911**
>
> **Platysilurus sp.**
>
> Figure 13.

**Figure 13:**
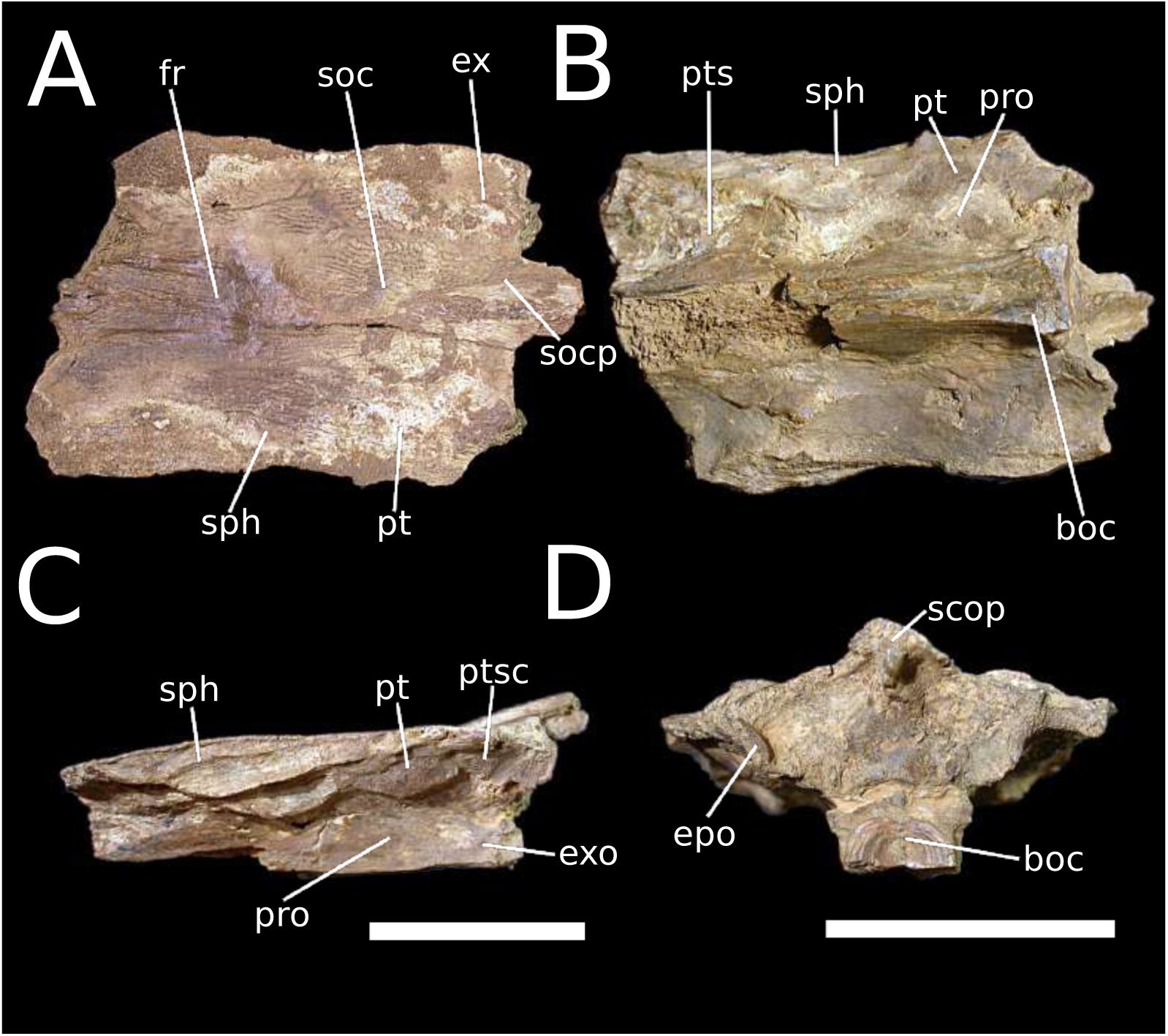
Partial neurocranium of *Platysilurus* sp., MUN 37605. A-D) Dorsal, ventral, lateral, and posterior views respectively. Scale bars equal 20 mm; the scale bar in C applies to A-C while the bar in D applies only to such section.

**Provenance**—Ware formation, Locality 470062, MUN 37605, partial neurocranium.

**Description**—Specimen preserving several cranial bones and surface ornament. Tridimensional structure compromised by squashing due to diagenesis, although most bones remain spatially in place. Sediment and iron cover tend to obscure most sutures between bones, making difficult to trace specific bone outlines. Dorsal bones (frontal, sphenotic, pterotic, supraoccipital, extrascapular) with ornament consisting of parallel and radiating ridges with origin foci in supraoccipital and pterotics; ridges sometimes undulating, often interleaved with scattered tubercles. Presence of evident depression on posteriormost region of frontals, anterior to suture with supraoccipital. Dorsal surface noticeably concave between level of pterotics to anteriormost preserved frontals, then strongly convex from supraoccipital ornament focus to preserved posterior limit of supraoccipital process. Absence of median longitudinal sulcus along supraoccipital process.

Frontal incomplete and preserving only about posterior half of original extent, showing medial depression posterior to region of anterior cranial fontanelle, surface covered with parallel, divergent, straight ridges; frontal in contact with supraoccipital and sphenotic, suture between frontal and supraoccipital strongly interdigitate, suture with sphenotic smooth and clean to slightly curve in some regions, suture with contralateral frontal mainly smooth with some degree of interdigitation at level of frontal depression posterior to anterior cranial fontanelle. Sphenotic elongate and renoind in outline, almost completely preserved in dorsal view; sphenotic in contact with frontal, supraoccipital, pterotic, pterosphenoid, and prootic; mesial suture with frontal as described above, posterolateral suture with pterotic nearly straight and smooth dorsally and interdigitate ventrolaterally, mesial suture with supraoccipital finely interdigitate, ventral suture with pterosphenoid concealed under sediment and bone fragments, ventral suture with prootic obscured due to lateral displacement of ventral wing of sphenotic during diagenesis, apparently smooth and straight. Lateral surface present yet squashed, although preserving spatial relationship to neighboring bones of otic region and some sutures. Hyomandibular articular facet elongate, concave, smooth, spanning sphenotic and anterior half of pterotic. Pterotic ovoid in dorsal view with posteromedian concavity for contact with exstrascapular; pterotic in contact with sphenotic, supraoccipital, extrascapular, prootic, postemporo-supracleithrum, and exoccipital; anteromesial suture with sphenotic as already described above, mesial suture with supraoccipital finely interdigitate, posterolateral suture with postemporo-supracleithrum and exoccipital obscured by fractures and sediment, ventral suture with prootic interdigitate although region of suture is fractured. Extrascapular barely visible in posterior view, strongly covered by sediment. Supraoccipital well preserved, polygonal in outline with posterior prominent supraoccipital process. Sutures as already described with adjacent bones.

Pterosphenoid poorly preserved, only posterior portion in contact with sphenotic and prootic present although very fractured and covered with sediment. Prootic in contact with pterosphenoid, sphenotic, pterotic, postemporo-supracleithrum, basioccipital, and exoccipital. Sutures with adjacent bones mostly obscured by fractures, however, preserved outlines suggest interdigitate sutures at least with basioccipital, exoccipital and sphenotic. Exoccipital in contact with prootic and basioccipital. Outlines poorly preserved, although interdigitate sutures are present with adjacent bones. Basioccipital poorly preserved, fractured, missing basal half, preserving articular facet for weberian apparatus in posterior view. Sutures with adjacent bones as already described above. Epioccipital poorly preserved, only part of bone exposed out of strong sediment cover, therefore obscuring sutures.

**Remarks**—Most of the fossil occurrences of the Ware formation so far are represented by very fragmentary specimens. Two notable exceptions are the weberian apparatus of *Brachyplatystoma* cf. *vaillantii* described by Aguilera et al. (2013) and the neurocranium of *Platysilurus* sp. herein described. The fossil record of the genus was until now restricted to the Urumaco and Rio Yuca formations in Venezuela (Sabaj Pérez et al., 2007; Lundberg et al., 2010; Rincón et al., 2016). The latter records occurrences are interesting because they are represented by the same anatomical region of the head as the specimen herein described; all of these specimens are also only identified to genus level and therefore its specific identity and affinities await re-study. The genus *Platysilurus* includes small to medium-sized piscivorous catfishes restricted to the Maracaibo drainage (*P. malarmo*), and Orinoco-Amazonas drainages (*P. mucosus*); it is readily distinguishable from other pimelodid genera by the highly raised supraoccipital process, a feature herein verified in museum specimens for the extant species, evident in the figures of Venezuelan fossil specimens (Sabaj Pérez et al., 2007; Rincón et al., 2016), and already suggested by Sabaj Pérez et al. (2007) as diagnostic for the genus. Despite the completeness of the material, family-scale comparative anatomy is still needed in order to determine whether the preserved characters are taxonomically informative. Both extant species show a prominent longitudinal sulcus along the supraoccipital process. *Platysilurus mucosus* shows continuity between interfrontal concatity and the longitudinal supraoccipital sulcus, in contrast with the specimen herein described where such connection is missing as well as the supraoccipital sulcus. *Platysilurus malarmo* has a nearly flat surface between the eyes anteriorly to before supraoccipital process, while the fossil specimen herein studied has a concave surface. Our fossil specimen is overall more similar to *P. malarmo* than to *P. mucosus*, despite the clear differences with either species already mentioned; this suggests that the fossil specimen might represent an extinct, undescribed species. Further detailed anatomical study of the congener *P. malarmo* is however needed before claiming any concise specific assignment for the fossil *Platysilurus* of the Ware formation. Specimens from the Urumaco and Rio Yuca formations are overall similar to the *Platysilurus* from the Ware formation; however, it was impossible to examine any of these specimens in order to decide whether they represent only one or different taxa. Given the cis- and trans-Andean distributions of both extant species, this occurrence may either support the presence or absence of drainage connections until its phylogenetic position in the family is better understood.

> **Genus *Zungaro* Bleeker, 1858**
>
> ***Zungaro* sp.**
>
> Figure 14.

**Figure 14:**
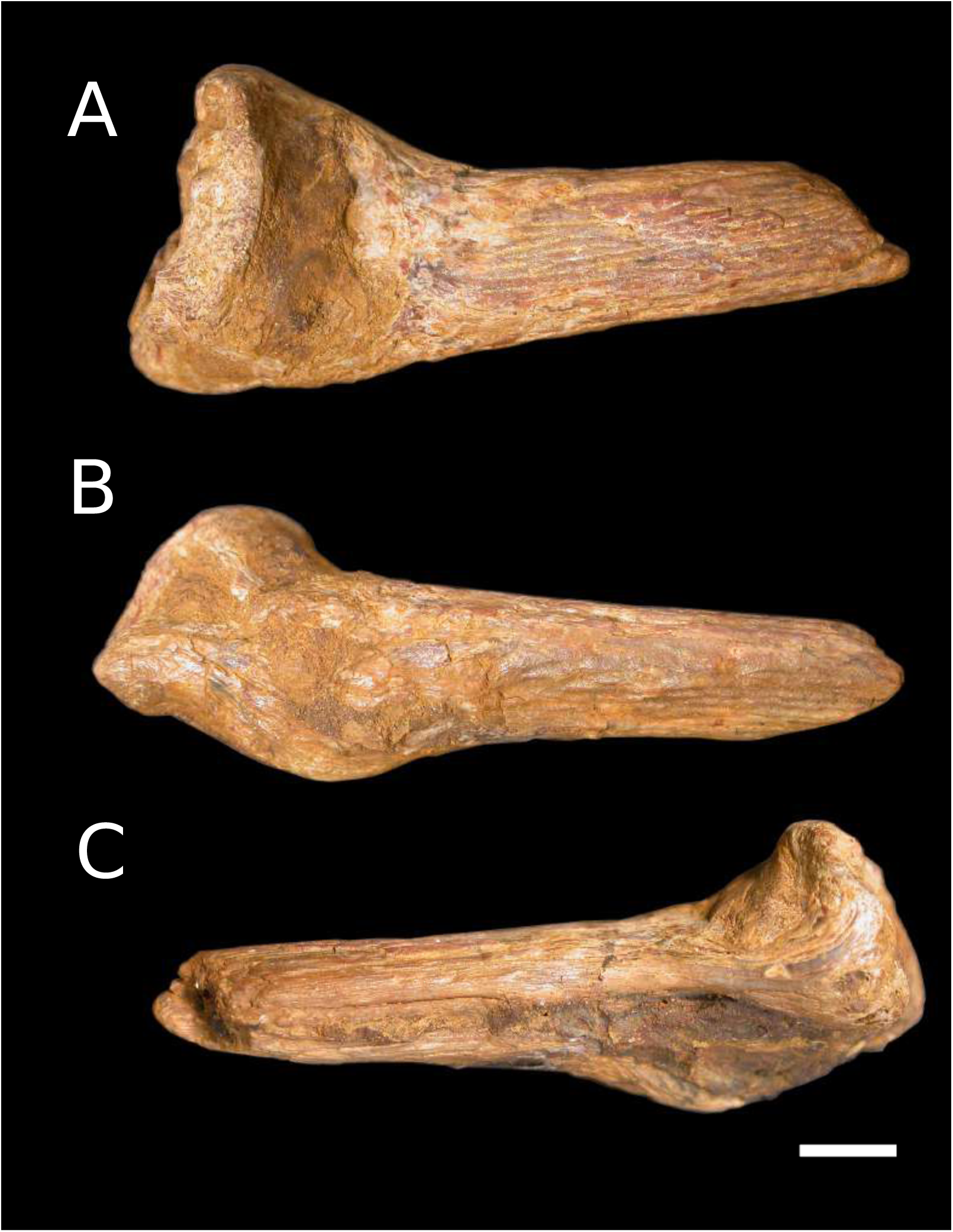
Left pectoral-fin spine of *Zungaro* sp., MUN 34483. A-C Dorsal, anterior, and posterior views respectively. Scale bar equals 5 mm.

**Provenance**—Ware formation, Locality 470060, MUN 34483, left partial pectoral spine preserving part of the spine base and proximal 1/4 of the shaft.

**Description**—Spine base preserving dorsal process with wide articular facet, otherwise strongly eroded, base of spine not preserving both proximal articular facet and proximal process. Anterior ornament absent; posterior ornament consisting of straight spinules; dorsal ornament consisting of smooth ridges, widest restricted to lateral 2/3 of shaft surface, additional narrow ones restricted to 1/3 mesial surface of shaft surface, oblique sulcus separating both kinds of surface ornaments.

**Remarks**—The genus *Zungaro* comprises *Z. jahu* and *Z. zungaro*, the former distributed in the Paraná-Paraguay drainage, and the latter in the Amazon and Orinoco drainages (Boni et al., 2011; Dagosta and de Pinna, 2019), therefore being one more representative of the Siluriformes restricted to cis-Andean South America. The fossil record of the genus includes a partial mesethmoid from the Rio Acre fossil fauna of Madre de Dios, Perú (LACM 128395, Lundberg et al., 2010). The present occurrence in the Guajira Peninsula represents the first trans-Andean record of the genus, a region outside of its current distribution. The combination of absent anterior ornamentation, and two regions with differential ridge thickness on the dorsal ornament of the pectoral-fin spine are diagnostic for the genus among Neotropical Siluriforms. *Zungaro* is a migratory species that occupies deep portions of lotic environments (Agostinho et al., 2003).

### 4.2 Faunal similarity

The freshwater fossil fish faunas herein studied vary considerably in richness of genera. The Urumaco fauna is the richest with 15 genera, followed by La Victoria and Villavieja with 14, while Loyola-Mangan, Sincelejo and Rio Yuca are the poorest with two genera each. The rest of the fossil assemblages vary from four to 13 genera. The Ware fauna scores seven out of the 17 fossil assembalges herein studied, with a comparatively large number of taxa (nine genera). According to the Bray-Curtis dissimilarity index, the Ware fauna is the most dissimilar of all the assemblages, most similar to Urumaco and a cluster of assemblages Austral, Amazonian, Orinocoan, and trans-Andean assemblages (Figure 15). This high degree of dissimilarity between Ware and other Neogene faunas of South America, either cis- or trans-Andean, is expected given that several genera are only known from the this fossil locality (e.g., *Hemidoras*, *Serrasalmus*, *Trachelyopterichthys*) while other taxa were shared with few other fossil localities (e.g., *Brachyplatystoma* and *Platysilurus*). The Loyola-Mangan and Solimões-Pebas faunas cluster together as one of the most dissimilar clusters in the set.

**Figure 15:**
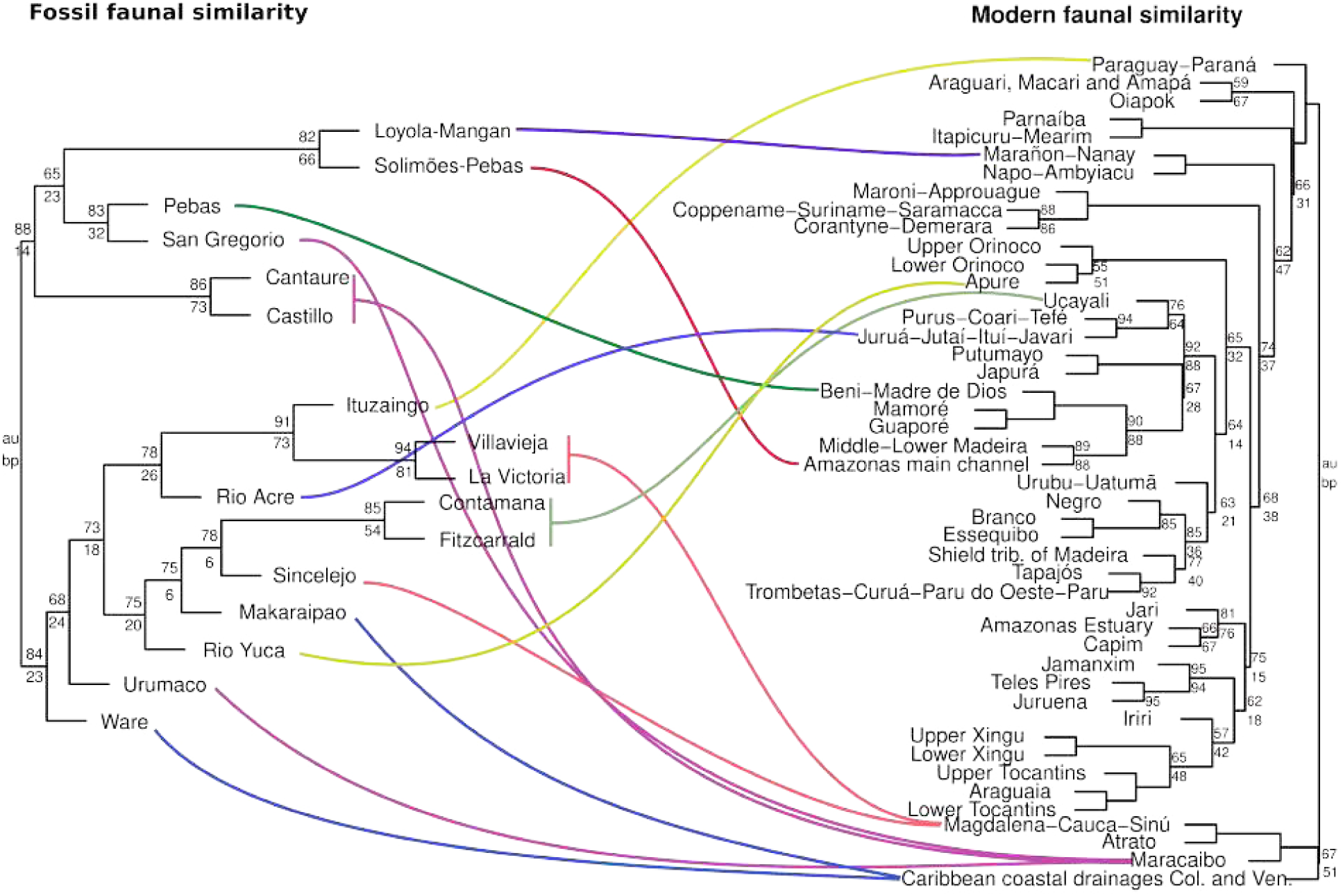
Faunal dissimilarity for Neogene freshwater fish faunas in South America using the Binary method. Colored lines match the fossil localities with their corresponding modern drainage. Colors are chosen only as visual aids to match recent and fossil assemblages. Support values above and below each node are approximately unbiased (AU) and bootstrap (BT) nodal support val-ues, respectively; only values lower than 95 are shown. The coastal drainages of Colombia (Col.) and Venezuela (Ven.) are small and numerous and drain the Andes into the Caribbean.

One of the most important features highlighted by this analysis is the lack of a clear correspondence between cis- and trans-Andean drainages which emerges when analysing the modern assemblages, where Magdalena-Cauca-Sinú, Atrato, Maracaibo, and Coastal drainages of Colombia and Venezuela group together in an exclusively trans-Andean, whereas the fossil assemblages do not show such association. The pattern of similarity does not seem to correlate with geographic proximity as was found by Ballen et al. (2021). However, it was not possible to test numerically the possible relationship between age and dissmilarity because the Pebas fossil assemblage (e.g., Solimões-Pebas, or Pebas itself) includes collections of faunas of different ages, spanning most of the Miocene (Hoorn et al., 2022).

## 5 Discussion

### 5.1 Informativeness of fragmentary specimens and taxonomic accuracy

Further field prospection will be necessary in order to expand our knowledge on the freshwater fishes of the Sincelejo formation, specially focusing on the facies with concretions where fossil vertebrates seem to be better represented (Figure 2C). Despite several field seasons and collection efforts throughout several years, few specimens of any vertebrate group have been recovered in the vicinity of Corozal when compared to rich fossil faunas such as La Venta and Ware, which suggests that recovery rate per rock volume tends to be low. Although original low concentration of remains in the sedimentary record is a possibility, other reasons such as low rock are exposure, fewer exposures with horizontal continuity, and vegetation cover might account for the poor recovery of specimens from Corozal when compared to the fossil record of the Ware and La Venta.

Exposures of the Ware formation, although regionally restricted, are locally well developed and devoid of vegetation cover as is the rule in the desertic Guajira Peninsula. Although numerous fossil specimens have been recovered in this unit, specially in the locality of the stratotype section, they suffer from excessive erosion and fragmentation, greatly reducing preservation of diagnostic features in fossil specimens. This explains why despite having numerous bone fragments available for study, only a portion have shown diagnostic features that allow to identify them to genus level.

### 5.2 New records

The present occurrences from the Sincelejo formation represent the first record of freshwater fishes from the Pliocene of the Magdalena-Cauca drainage (Figure 16). This component of the extinct continental diversity in trans-Andean Northern South America has long been neglected despite being known from a number of stratigraphic units in the Magdalena-Cauca drainage since the first half of the XX century (e.g., the Coyaima, La Venta, Carmen de Apicalá faunas; Stirton, 1953, pp. 610, 612, 616). Among upper Magdalena faunas historically known to contain freshwater fishes, only the La Venta fauna has been extensively studied (Lundberg et al., 1986; Lundberg and Chernoff, 1992; Lundberg, 1997, 2005; Lundberg et al., 2010; Ballen and Moreno-Bernal, 2019), with other records restricted to order-level identification or simply indeterminate fish remains. Herein, one remain was identified to genus level and another to family.

**Figure 16:**
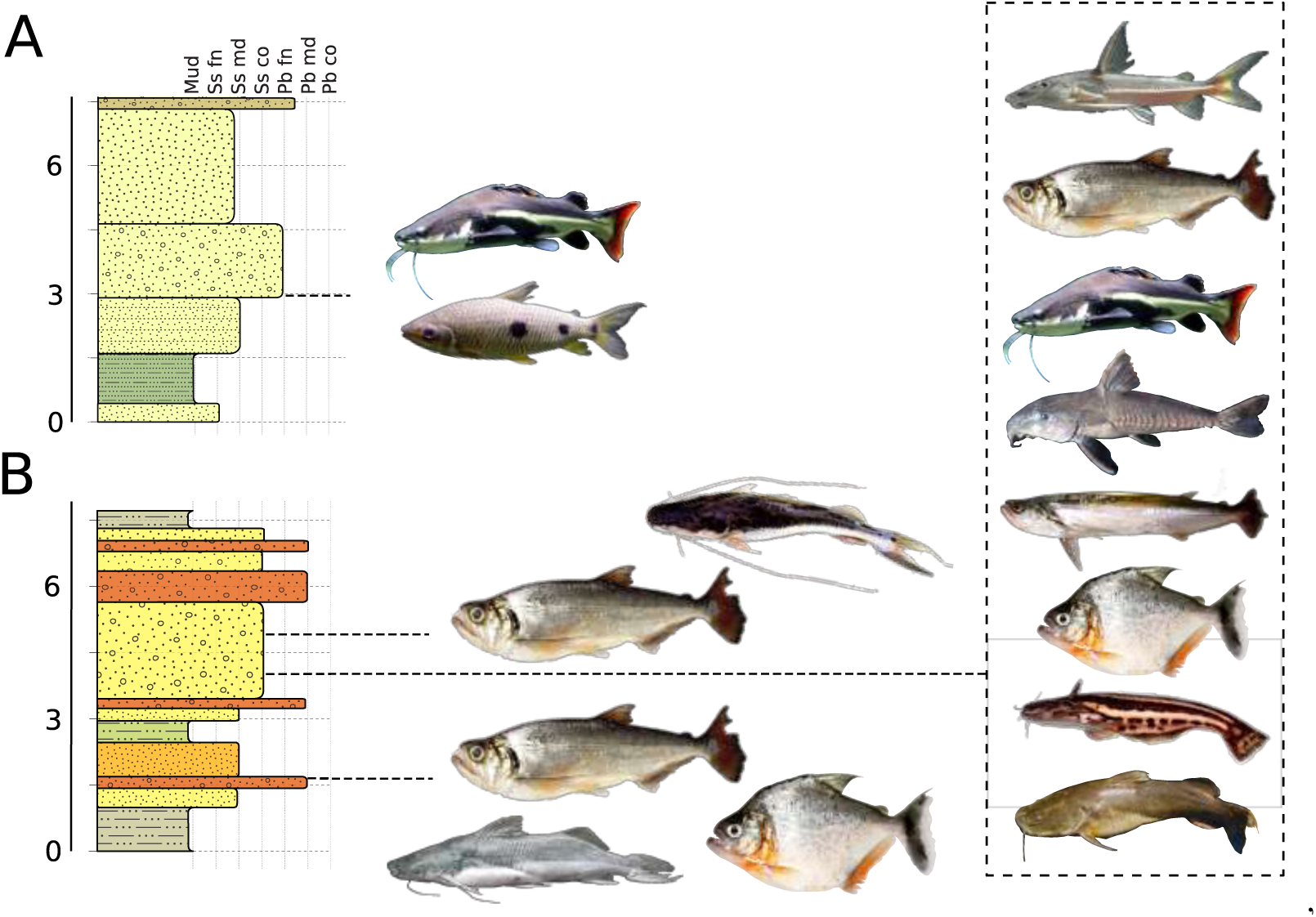
Stratigraphic distribution of the taxa recovered in this study. A) San Francisco farm section with *Leporinus* or *Hypomasticus* and *Phractocephalus* sp. B) Ware formation stratotype with *Brachyplatystoma* cf. *vaillantii*, *Hydrolycus* spp., and *Serrasalmus* at 1.7m; *Hemidoras* sp., *Hydrolycus* spp., *Phractocephalus* sp., cf. *Oxydoras* sp., cf. *Rhaphiodon* sp., *Serrasalmus* sp., *Trachelyopterichthys* sp., and *Zungaro* sp. at 4m; *Hydrolycus* spp. and *Platysilurus* sp. at 5m. Staff scales in stratigraphic meters. Vertical guides are granulometry; Mud = mud, Ss = sand, Pb = pebbles, fn = fine, md = medium, co = coarse. Stratigraphic columns as in Figure 3. Sources of the pictures: Cliff on wikicommons (*Phractocephalus hemioliopterus*), ANSP/G.W. Saul (*Zungaro zungaro*), Jonathan Armbruster (*Oxydoras niger*), Citron on wikicommons (*Leporinus friderici*), Clinton & Charles Robertson (*Hydrolycus tatauaia*, *Serrasalmus* sp., *Rhaphiodon vulpinus*, *Trachelyopterichthys taeniatus*), Mark Sabaj (*Brachyplatystoma vaillantii*, *Hemidoras morrisi*), and Galvis et al. (1997, *Platysilurus malarmo*).

The Ware fauna is by far the most rich of both faunas herein studied with ten recognized taxa, mostly to genus level (Table 2, Figure 16). As mentioned, the genera *Hemidoras*, *Serrasalmus*, and *Trachelyopterichthys* are new records for the fossil freshwater fishes of South America. The genera *Platysilurus*, *Phractocephalus*, and *Zungaro* are already known from other fossil faunas of South America; however, all records herein reported are new for the trans-Andean region, and almost all being also the youngest of the fossil record of their respective groups (Lundberg et al., 2010). The only exception is the genus *Platysilurus*, whose occurrence in the Rio Yuca formation of Venezuela is also of Pliocene age (Bermúdez et al., 2015). The genus *Pygocentrus* in the La Venta and Contamana faunas are the first reliable occurrences of the genus in the fossil record.

**Table 2:**
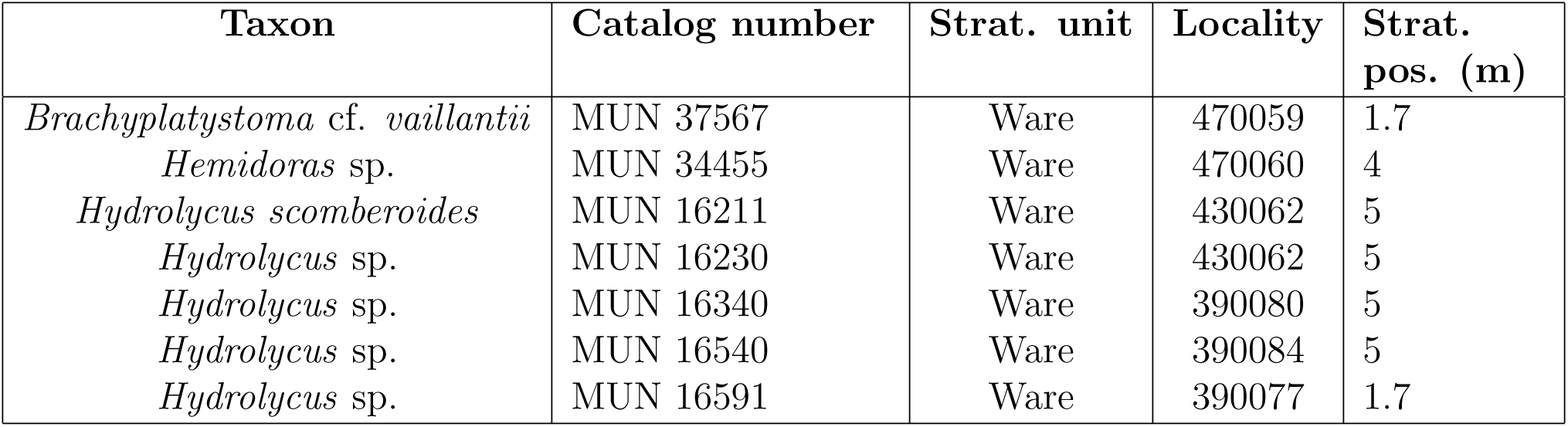
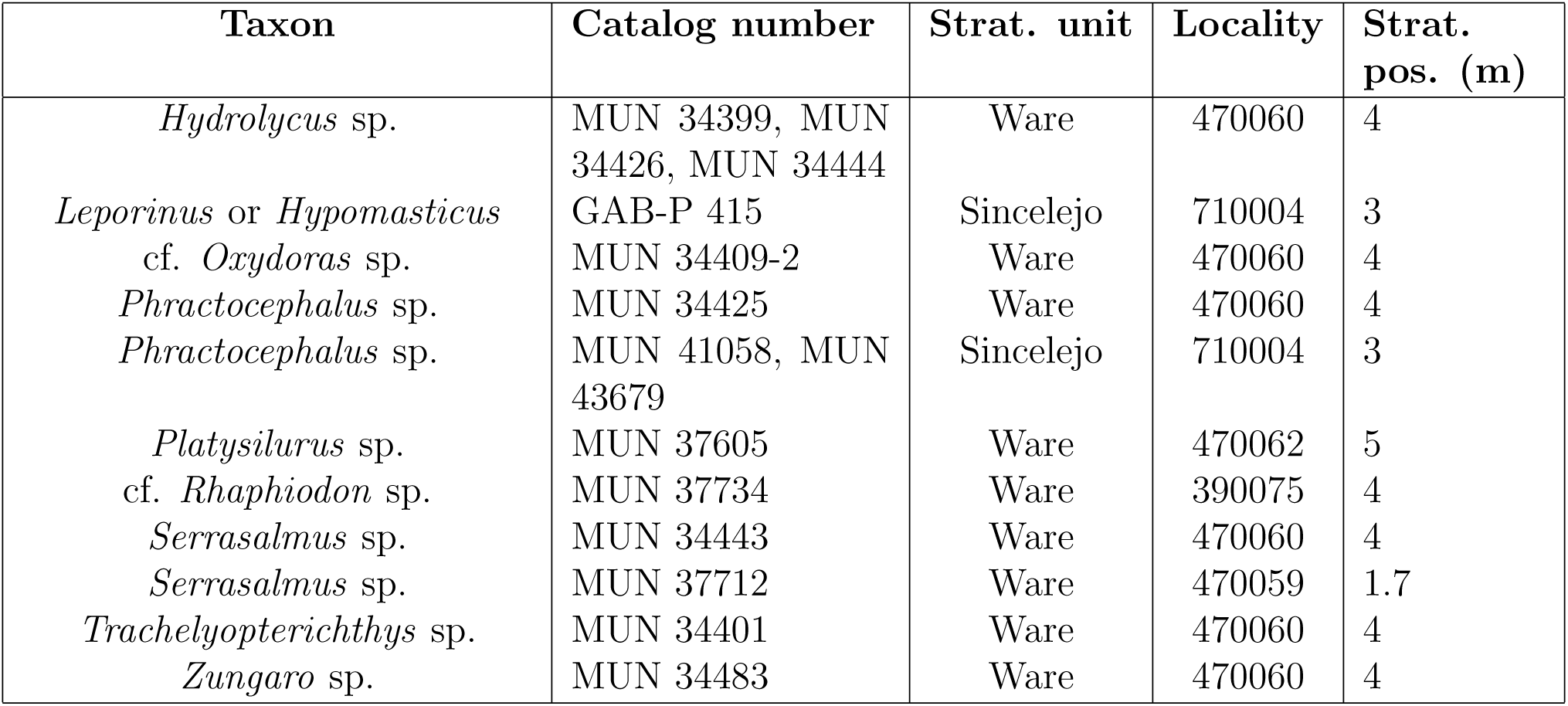
STRI localities from the Sincelejo and Ware formations referenced in the present study. Several localities refer to the same level and spatial point, being therefore synonyms, see Figure 3.

| Taxon | Catalog number | Strat. unit | Locality | Strat. pos. (m) |
| --- | --- | --- | --- | --- |
| *Brachyplatystoma* cf. *vaillantii* | MUN 37567 | Ware | 470059 | 1.7 |
| *Hemidoras* sp. | MUN 34455 | Ware | 470060 | 4 |
| *Hydrolycus scomberoides* | MUN 16211 | Ware | 430062 | 5 |
| *Hydrolycus* sp. | MUN 16230 | Ware | 430062 | 5 |
| *Hydrolycus* sp. | MUN 16340 | Ware | 390080 | 5 |
| *Hydrolycus* sp. | MUN 16540 | Ware | 390084 | 5 |
| *Hydrolycus* sp. | MUN 16591 | Ware | 390077 | 1.7 |

Table 2 – *Continued from previous page*
| **Taxon** | **Catalog number** | **Strat. unit** | **Locality** | **Strat.**<br>**pos. (m)** |
| --- | --- | --- | --- | --- |
| *Hydrolycus* sp. | MUN 34399, MUN<br>34426, MUN 34444 | Ware | 470060 | 4 |
| *Leporinus* or *Hypomasticus* | GAB-P 415 | Sincelejo | 710004 | 3 |
| cf. *Oxydoras* sp. | MUN 34409-2 | Ware | 470060 | 4 |
| *Phractocephalus* sp. | MUN 34425 | Ware | 470060 | 4 |
| *Phractocephalus* sp. | MUN 41058, MUN<br>43679 | Sincelejo | 710004 | 3 |
| *Platysilurus* sp. | MUN 37605 | Ware | 470062 | 5 |
| cf. *Rhaphiodon* sp. | MUN 37734 | Ware | 390075 | 4 |
| *Serrasalmus* sp. | MUN 34443 | Ware | 470060 | 4 |
| *Serrasalmus* sp. | MUN 37712 | Ware | 470059 | 1.7 |
| *Trachelyopterichthys* sp. | MUN 34401 | Ware | 470060 | 4 |
| *Zungaro* sp. | MUN 34483 | Ware | 470060 | 4 |

### 5.3 Palaeoenvironments

Fossil-bearing facies of both the Sincelejo and Ware formations share granulometric and sedimentary features typical of high-energy environments such as rivers, subject to high rates of transport and erosion. Although transport should have taken place in such a high-energy depositional environment, this process appears to have been only local in scale, as judging from the size class mixture of the collected remains (small fragments at the millimeter order of magnitude are mixed with centimeter-scale particles). Large-scale transport, on the other hand, would have generated a stronger size sorting that is not present in the fossiliferous facies of the Ware stratopype, where large-sized remains such as crocodylian bones are intermingled with small-sized fragments such as catfish spine fragments and piranha teeth, roughly spanning the cm to mm orders of magnitude. On the other hand, if large-scale transport was present and mixed with local supply of bone fragments, differential degrees of erosion would be present with small specimens better preserved than large-sized remains, the latter having traveled a longer erosive distance. Detailed taphonomic studies could shed light on the likely degree of transport and sorting in this fossil assemblage.

Some of the taxa recovered show certain degree of ecological specialization. For instance, large catfish species today (e.g., *Brachyplatystoma*, *Phractocephalus*, *Platysilurus*, and *Zungaro*) are known to undergo seasonal reproductive migrations in large rivers, therefore requiring large drainages (Carolsfeld et al., 2003). This combination of taxa strongly suggest that the Ware formation is composed by sediments of a medium to large-sized river that was part of a large drainage where these taxa could migrate. Cynodontid fishes (*Hydrolycus* and *Rhaphiodon*) apparently also take part in migrations, although not to the same distances as pimelodid catfishes.

Most of the taxa preserved in the Ware and Sincelejo formations are of the piscivorous– carnivorous trophic guild, with a notable absence of herbivorous especialists such as pacus and common omnivores such as characids; this can be explained by the fact that these high-order nodes in trophic networks tend to be the largest in body size in Neotropical environments, and facilitates its recovery in fossil samples while more fragile and smaller fishes tend to be less represented or even absent at all. Pacu teeth (e.g., see Ballen et al., 2021) are very resistant and their remains are expected to be recovered in future collection efforts as they are a prominent component in most South American Neogene faunas. A minor proportion tends to be more directed towards detritivory (e.g., the Doradidae).

Although the Characidae is a prominent, very rich, abundant and important component of the biomass in fish communities in the Neotropics, they are rarely represented in fossil samples and are generally restricted to isolated teeth.Abrasion easily destroys the laminar and fragile bones of these fishes. This also explains the fact that piranhas are only known from isolated teeth since their bones are as fragile as those of most Characiformes. Such preservation bias explains the general lack of representativeness of this order in the fossil record in South America (Lundberg et al., 2010).

The Ware fish fauna requires permanent medium size to large rivers that highly contrast with desertic landscape of the northern Guajira nowadays undercoring that the region has gone through an intense phase of desertification over the past few million years as other analysis also have indicated (Jaramillo et al., 2020).

The dissimilarity analysis recovered a pattern consistent with previous analyses prior to the study of the Ware fauna (Figure 15). The main difference relative to previous analyses (figure 6 in Ballen et al., 2021) is the position of the Ware fauna, at the base of the other Neogene fossil faunas. As was found by these authors, this pattern is expected given the amount of taxa that are unique to the Ware fauna, as well as the few shared components with other fossil faunas. The lack of correlation with geographic proximity is a pattern already recovered in previous studies using mammal faunas (Carrillo et al., 2015), and freshwater fishes (Ballen et al., 2021). This pattern may be affected by an insufficient knowledge of the Peruvian faunas, or perhaps highlight ecological heterogeneity among sites. Until the peruvian fish faunas are studied in detail, we might not be able to distinguish between the two alternatives.

### 5.4 Palaeogeography

The rich fossil assemblage herein studied shows a consistent pattern of association with cis-Andean groups. As already mentioned, both taxa from the Sincelejo fauna are currently restricted to drainages west to the Andes (*Leporinus* or *Hypomasticus*, and *Phractocephalus*), while the phylogenetic position of the latter reinforces its palaeogeographic relevance. The summed fossil occurrences of the Sincelejo and Ware formations suggest that there was a drainage connection between the Magdalena-Cauca drainage and the Amazon-Orinoco drainages by the Pliocene, or alternatively, that these taxa survived in the isolated drainages until Pliocene times (Lundberg and Chernoff, 1992).

The Ware fauna is composed of taxa that are consistently part of groups restricted to the drainages east to the Andes (*Brachyplatystoma*, *Hemidoras*, *Hydrolycus*, *Phractocephalus*, *Serrasalmus*, *Trachelyopterichthys*, and *Zungaro*). The doubtful occurrence of *Rhaphiodon* also adds to this pattern, because their family is entirely restricted to cis-Andean drainages. The genus *Platysilurus* provides ambiguous palaeogeographic information given that one of its extant species is cis-Andean while the other is found in the Maracaibo drainage (trans-Andean). *Platysilurus* from Ware and potentially those from the Rio Yuca and Urumaco formations in Venezuela do not seem conspecific with *Platysilurus malarmo*, the trans-Andean species. The only occurrence confidently identified to species level, *Hydrolycus scomberoides*, is a component restricted to the Amazon drainage, today absent in the Orinoco drainage, therefore reinforcing the association suggested by other taxa; it is intriguing that this species is currently absent in the Orinoco drainage while present in northern Colombia by the Pliocene. This indicates a case of regional extirpation.

The strong resemblance between the Sincelejo and Ware faunas on the one hand, and cis-Andean freshwater fish communities on the other, is not an isolated pattern. It conforms with the same composition found in the La Venta fauna, where almost all of the taxa in the fossil record are nowadays restricted to the Amazon drainage (Lundberg, 1997, 2005; Ballen and Moreno-Bernal, 2019). Present results would support the model of drainage connection to the mid- to late Miocene by Lundberg (1997) for the La Venta fauna. The present data suggest that the connection could be maintaned until the Pliocene, suggesting that some kind of drainage connection was still present at that period across the Cordillera Oriental of Colombia and/or the Merida Andes in Venezuela. A possible geological pattern explaining this persistence is the late uplift of the Garzón massif in the southern Cordillera Oriental during the Pliocene, creating room for drainage connection between the upper Magdalena and the Amazon-Orinoco drainages (Anderson et al., 2016; Saeid et al., 2017) (Figure 17). This kind of connection would explain both the fish composition and the required drainage properties such as a long course and connection to cis-Andean drainages. Alternative mechanisms such as connections across the Merida Andes in Venezuela have also been suggested in the literature, although they seem less likely as they require extensive uplift in a large area during the past million years (Audemard and Audemard, 2002; Audemard, 2003). The latter model has also been challenged in the literature because the present-day elevation of the Merida Andes is thought to have been reached during the late Miocene to Pliocene (Bermúdez et al., 2015). An alternative scenario could be that all modern cis-Andean lineages continue in drainages west of the Andes over millions of years following the separation of both regions, and its trans-Andean extirpation occurred at some point during the late Pliocene or Pleistocene due to the loss or decrease of river drainages due to aridification (e.g. Ware, and La Venta). Further combination of fossil information, molecular and morphological data, tectonic models, and complex probabilistic modeling has the potential to provide a better understanding of the process of mountain building, its associated drainage evolution, and ultimately the evolution of the neotropical biota.

**Figure 17:**
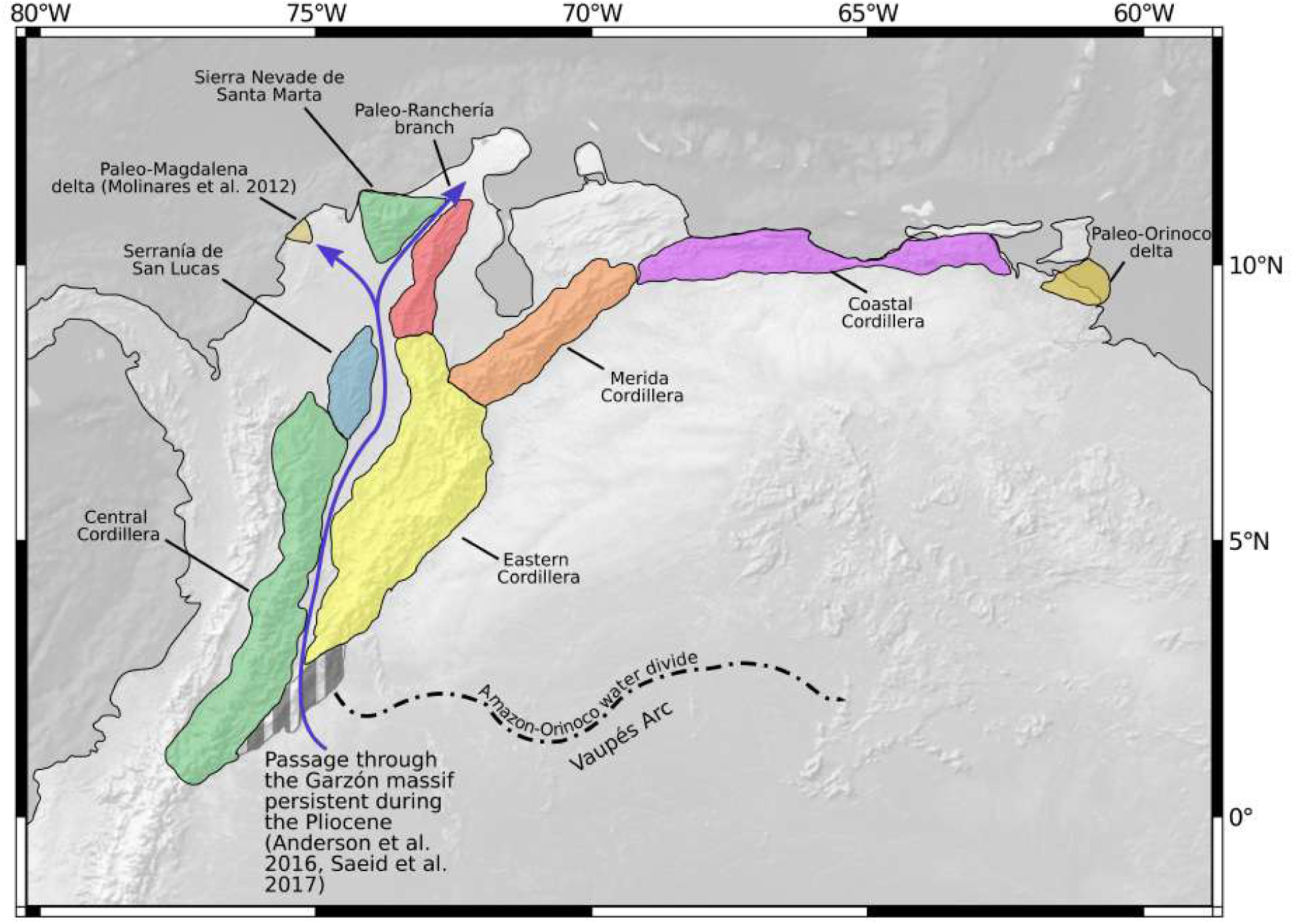
Palaeogeography of northern South America showing a possible palaeodrainage connection between drainages east and west of the Andes through the Garzón area which was likely not uplifted yet during the Pliocene (ca. 2–4 Ma).

## 6 Acknowledgments

This study was funded through FAPESP grants and fellowships to GAB (2014/11558-5, 2016/02253-1, and 2023/07838-1), NSF grant EAR-0957679, a Böhlke Travel Grant by the Academy of Natural Sciences of Drexel University, the National Geographic Society, and the Smithsonian Tropical Research Institute. We thank the MAPUKA museum (Universidad del Norte) for granting access to fossil specimens. John G. Lundberg (ANSP), the late Rich Vari (NMNH), the late Iván Mojica (ICNMHN), and Aléssio Datovo (MZUSP) are acknowledged for providing access to collections under their care. The Center for Tropical Paleoecology and Archaeology (Smithsonian Tropical Research Institute), the Academy of Natural Sciencies of Drexel University, the Instituto de Ciencias Naturales (Universidad Nacional de Colombia), the Pontificia Universidad Javeriana of Bogotá, and the Museu de Zoologia da Universidade de São Paulo provided logistic support. Thanks to the comminities of Warpana, Patajau, Aulechit, Nazareth, Wososopo, Sillamana, Paraguachón, Flor de la Guajira, and Ipapura. Thanks to the Colombian National Police (Castilletes base) and the Colombian Army (La Flor de la Guajira and Cerro de la Teta). Part of the analyses were run on a server at the IBB/UNESP Botucatu (FAPESP process 2014/26508-3), Claudio Oliveira and Fábio Roxo are acknowledged for granting access to computational infrastructure at UNESP Botucatu. Gustavo Burin, Flavio Lima, Fernando Dagosta, Victor Tagliacollo, Max Langer, Naércio Menezes, and Murilo Pastana are acknowledged for fruitful discussions during the preparation of the manuscript and final phases of analysis. We thank Federico Moreno, Maria Camila Vallejo, Orangel Aguilera, and John Lundberg for discussions on the paleogeography of northern South America.

