## Supplementary material for "Fossil Freshwater Fishes from the Pliocene of northern Colombia and the Palaeogeography of northern South America"

December 18, 2024

#### 1 Appendix: Material examined

**ACTINOPTERYGII**
**CHARACIFORMES**

##### ACESTRORHYNCHIDAE

*Acestrorhynchus falcirostris*: MZUSP 92991 (1 out of 2, DS). *Acestrorhynchus heterolepis*: MZUSP 82896 (1, DS). *Acestrorhynchus lacustris*: MZUSP 83376 (1, DS). *Gilbertolus alatus*: ICNMHN 16753 (16, 99.9–120.0 mm SL), BMNH 1924.3.3.46–48 (1, CS). *Gilbertolus atratoensis*: MZUSP 10663 (1, CS). *Roestes molossus*: INPA 11068 (1, CS), MZUSP 52063 (1, CS). *Roestes ogilviei*: ICNMHN 144699 (1, 120.0 mm SL).

**CHARACIDAE**

*Cynopotamus atratoensis*: USNM 310494 (2, 142.1–144.3 mm SL).

**CYNODONTIDAE**

*Cynodon gibbus*: MZUSP 15641 (1, CS), USNM 222848 (1, 205.0 mm SL), USNM 257564 (1, 197.0 mm SL), USNM 305368 (2, 167.0–240.0 mm SL), USNM 403923 (1, 250 mm SL). *Cynodon meionactis*: MZUSP 99683 (1, 151.9 mm SL). *Cynodon* sp.: MZUSP 91687 (1, DS). *Cynodon septenarius*: MZUSP 92544 (3, 189.2–195.5 mm SL). *Hydrolycus armatus*: MZUSP 94477 (1, DS), 89507 (2, DS), 95879 (1, DS), USNM 403920 (1, 245.0 mm SL), USNM 403921 (1, 235.0 mm SL). *Hydrolycus scomberoides*: ANSP 159099 (1, DS), MZUSP 6971 (1, CS), 26177 (1, CS), 32093 (1, CS), 32616 (1 out of 8, 175.3 mm SL, dentary as DS preparation), USNM 403923 (1, 250.0 mm SL). *Hydrolycus tatauaia*: MZUSP 94086 (3 out of 4, DS), 32637 (1, CS), USNM 402147 (1, 200.0 mm SL). *"Hydrolycus" wallacei*: MZUSP 32634 (1, 269.0 mm SL, dentary as DS preparation), 32638 (1, CS). *Rhaphiodon vulpinus*: MZUSP 117050 (2, DS), 92008 (1, DS), USNM 52549 (4, 250.0–300.0 mm SL), USNM 126662 (5, 128.8–184.0 mm SL), USNM 222863 (1, 233.0 mm SL), USNM 310941 (2, 198.0–265.0 mm SL).

**SERRASALMIDAE**

*Acnodon* sp.: MZUSP 20361. *Catoprion mento* MZUSP 20246 (10) *Metynnis fasciatus*: MZUSP 20555. *Mylesinus schomburgkii*: MZUSP 101455. *Myleus setiger*: MZUSP 15673. *Myleus* sp.: MZUSP 91685. *Myloplus rhomboidalis*: MZUSP 102136. *Myloplus schomburgkii*: MZUSP 97639. *Myloplus torquatus*: MZUSP 95388. *Mylossoma acanthogaster*: IAvH-P 15900. *Mylossoma duriventre*: MZUSP 89523. *Mylossoma paraguayensis* MZUSP 895232. *Piaractus brachypomus*: MZUSP 20376. *Piaractus mesopotamicus*: MZUSP 89508. *Pygocentrus nattereri*: MZUSP 89511. *Pygopristis denticulata*: MZUSP 57580. *Tometes kranponhah*: MZUSP 110948. *Utiaritichthys sennaebragai*: MZUSP 93692

#### SILURIFORMES

**ARIIDAE**

*Sciades proops*: MZUSP 52842.

**ASTROBLEPIDAE**

*Astroblepus* sp.: MZUSP 22326.

**ASPREDINIDAE**

*Bunocephalus coracoideus*: MZUSP 103254.

**AUCHENIPTERIDAE**

*Ageneiosus pardalis*: ICNMHN 10014. *Liosomadoras morrowi*: ICNMHN 14026. *Liosomadoras oncinus*: MZUSP 105828. *Tetranematichthys quadrifilis*: ICNMHN 12385. *Tocantinsia piresi*: MZUSP 100031, 98301, 103244, 103415. *Trachelyopterichthys anduzei*: ICNMHN 16904. *Trachelyopterichthys taeniatus*: MZUSP 6642. *Trachelyopterus striatulus*: MZUSP 86012. *Trachelyopterus insignis*: ICNMHN uncat. *Trachycorystes trachycorystes*: ICNMHN 2761, MZUSP 104547, 111057. *Trachycorystes porosus*: MZUSP 101876. *Trachycorystes* sp.: MZUSP 91659.

**BAGRIDAE**

*Mystus gulio*: MZUSP 63581.

**CALLICHTHYIDAE**

*Aspidoras albater*: MZUSP 28599, 28, 14.59–39.76; 50157, 1, cs. *Aspidoras microgalaeus*: MZUSP 86842, 1, cs. *Callichthys callichthys*: MZUSP 43599, 1, 121.12; MZUSP 93043, 8,

47.68–72.60; MZUSP 109087, 2, 116.71–145.12; MZUSP 84199, 2, cs. *Callichthys serralabiatum*: MZUSP 93168, 1, 104.65. *Corydoras araguaiaensis*: MZUSP 86269, 1, cs. *Corydoras ehrhardti*: MZUSP 81572. *Corydoras* cf. *guianensis*: MZUSP 107151, 2, cs. *Corydoras splendens*: MZUSP 89377, 5, 47.16–57.47. *Dianema* sp.: MZUSP 30862, 2, cs. *Hoplosternum littorale*: MZUSP 85987, 1, 84.91; MZUSP 94658, 1, 95.46; MZUSP 117107, 1, cs. *Lepthoplosternum pectorale*: MZUSP 83608, 2, cs; MZUSP 112403, 1, 57.13. *Megalechis thorocata*: MZUSP 25451, 2, cs. *Scleromystax barbatus*: MZUSP 37723, 2, cs.

**CETOPSIDAE**

*Cetopsis gobioides*: MZUSP 99279.

**CLARIIDAE**

*Clariallabes longicauda*: BMNH 1937.12.12:22. *Clariallabes melas*: MRAC 51599-611. *Dinotopterus cunningtoni*: UMMZ 199855. *Clarias* sp.: MZUSP 91656.

**CLAROTEIDAE**

*Chrysichthys auratus*: BMNH 1907.13.2:2082-7. *Phyllonemus filinemus*: MRAC 90257.

**DIPLOMYSTIDAE**

*Diplomystes camposensis*: MZUSP 88533. *Diplomystes viedmensis*: ANSP 192904.

**CRANOGLANIDIDAE**

*Cranoglanis bouderius*: ANSP 164978 (CT available at http://catfishbone.acnatsci.org/Cranoglanididae/Cranoglanis/bouderius/catscan.html).

**DORADIDAE**

*Acanthodoras cataphractus*: MZUSP 31098 (1). *Acanthodoras spinosissimus*: ICNMHN 8266, MZUSP 112857, MZUSP 29660 (1). *Agamyxis albomaculatus*: ICNMHN 1319, MZUSP 88607. *Agamyxis pectinifrons*: MZUSP 51777 (1). *Amblydoras affinis*: ICNMHN 8287, MZUSP 31699 (1). *Amblydoras bolivarensis*: MZUSP 88610. *Anadoras grypus*: ICNMHN 8274 MZUSP 86818, MZUSP 36119 (7). *Anadoras weddellii*: MZUSP 31701 (1). *Anduzedoras oxyrhynchus*: MZUSP 29025 (2). *Astrodoras asterifrons*: MZUSP 29049, MZUSP 111428 (2). *Centrodoras brachiatus*: MZUSP 103891 (2). *Centrodoras hasemani*: MZUSP 91675 (1). *Centrochir crocodili*: ICNMHN 5698. *Doras carinatus*: MZUSP 108947 (1). *Doras higuchii*: MZUSP 101627 (1). *Doras zuanoni*: MZUSP 110808 (1). *Franciscodoras marmoratus*: MZUSP 2201 (7). *Hassar affinis*: MZUSP 74809 (6). *Hassar gabiru*: MZUSP 111342 (10). *Hassar orestis*: MZUSP 22057 (1), MZUSP 82256 (1), MZUSP 21915 (1). *Hassar wilderi*: MZUSP 62998 (5). *Hemidoras morei*: MZUSP MZUSP 27844 (1). *Hemidoras morrisi*: MZUSP 82890 (4).x *Hemidoras stenopeltis*: ICNMHN 5936, MZUSP 42772 (4), MZUSP 7541 (4). *Hemidoras stuebelii*: ICNMHN 17104. *Hypodoras forficulatus*: ICNMHN 10143. *Kalyptodoras bahiensis*: MZUSP 100737 (4). *Leptodoras acipenserinus*: MZUSP 49774 (1). *Leptodoras cataniai*: MZUSP 82978 (2). *Leptodoras hasemani*: MZUSP 111855 (3). *Leptodoras juruensis*: MZUSP 82985 (1), MZUSP 82880 (2). *Leptodoras nelsoni*: ICNMHN 413. *Leptodoras oyakawai*: MZUSP 98216 (3). *Leptodoras praelongus*: MZUSP 31694 (1). *Leptodoras* sp.: ICNMHN 8422. *Lithodoras dorsalis*: MZUSP 46587 (1), MZUSP 74677 (4). *Megalodoras uranoscopus*: ICNMHN 6996, MZUSP 46007 (1), MZUSP 55838 (5). *Merodoras nheco*: MZUSP 109766 (1). *Nemadoras elongatus*: ICNMHN 8399, MZUSP 83211 (2). *Nemadoras hemipeltis*: MZUSP 83262 (1). *Nemadoras humeralis*: MZUSP 103873 (1). *Orinocodoras eigenmanni*: ICNMHN 8441, MZUSP 86807 (1). *Ossancora eigenmanni*: MZUSP 95024 (66), MZUSP 31709 (1). *Ossancora fimbriata*: MZUSP 56703 (39), MZUSP 52534 (3). *Ossancora punctata*: ICNMHN 16705, MZUSP 82894 (12). *Oxydoras niger*: IC-

NMHN 8445, MZUSP 56870 (1), MZUSP 86819 (3). *Oxydoras sifontesi*: MZUSP 76540 (1). *Physopixys lyra*: MZUSP 62709, MZUSP 121168 (6). *Physopyxis ananas*: MZUSP 117504 (9). *Platydoras armatulus*: ICNMHN 10025, MZUSP 90425 (1), MZUSP 116542 (5). *Platydoras costatus*: MZUSP 37008 (1). *Pterodoras granulosus*: ICNMHN 8300, MZUSP 91655, MZUSP 36100 (2). *Pterodoras rivasi*: MZUSP 88613 (1). *Rhinodoras armbrusteri*: MZUSP 125377 (1). *Rhinodoras dorbignyi*: MZUSP 106761 (3). *Rhinodoras thomersoni*: ICNMHN 2172. *Rhynchodoras woodsi*: MZUSP 86816 (1). *Scorpiodoras heckelii*: ICNMHN 12962, MZUSP 112942, MZUSP 85494 (1), MZUSP 36251 (1). *Tenellus leporhinus*: MZUSP 97520 (1). *Tenellus trimaculatus*: MZUSP 92344 (1). *Tenellus ternetzi*: MZUSP 55760 (3). *Trachydoras nattereri*: MZUSP 92372 (2). *Trachydoras paraguayensis*: MZUSP 62700 (2). *Trachydoras steindachneri*: MZUSP 49526 (5). *Wertheimeria maculata*: MZUSP 106759 (1).

**HEPTAPTERIDAE**

*Goeldiella eques*: MZUSP 33190. *Pimelodella chagresi*: ICNMHN 1623. *Rhamdia quelen*: MZUSP 102718.

**HETEROPNEUSTIDAE**

*Heteropneustes fossilis*: USNM 273737.

**LORICARIIDAE**

*Ancistrus centrolepis*: IAvH-P 10473, ICNMHN 104, 189, 1632, 3153. *Ancistrus martini*: ICNMHN 1206, 17647, 17648, 17653. *Ancistrus triradiatus*: ICNMHN 17649, 17654. *Baryancistrus niveatus*: MNRJ 19344. *Chaetostoma alternifasciatum*: ANSP 71711 (holotype, photograph, xray). *Chaetostoma anale*: ANSP 70525 (holotype), ICNMHN 13397, 17634. *Chaetostoma anomalum*: USNM 133135 (syntype, photograph, xray). *Chaetostoma breve*: BMNH 1898.11.4.33-36 (syntypes, photograph). *Chaetostoma brevilabiatum*: ICNMHN 6134 (holotype). *Chaetostoma carrioni*: BMNH 1933.5.29.1 (holotype, photograph). *Chaetostoma dorsale*: ICNMHN 1183, 3372, 3535, 7997, 8011, 8013, 8027, 8031, 17499, 17646, MLS 588, 747, 604. *Chaetostoma jegui*: INPA Uncatalogued (photograph). *Chaetostoma loborhynchos*: MUSM 20291 (photograph), CZUT-IC 5551, 5552. *Chaetostoma microps*: BMNH 1860.6.16.137-143 (syntypes, photograph). *Chaetostoma milesi*: ANSP 69330 (holotype), ICNMHN 10420, 15528, 16123, 16268, 16291, 16923, MLS 562. *Chaetostoma pearsei*: ICNMHN 10361. *Chaetostoma platyrhynchus*: ICNMHN 5488, 5492, 7971, 9417, 17624, 17625, 17626, 17628, 17629, 17630, 17631. *Chaetostoma sovichthys*: ICNMHN 2381, 16221, 16223, MLS 568, 590, 600, USNM 121053 (holotype, photograph, xray). *Chaetostoma* sp. "Perú": AUM 45597, 45634. *Chaetostoma tachiraense*: MLS 797, 799, 805, USNM 121052 (holotype, photograph, xray). *Chaetostoma vagum*: ANSP 70521 (holotype, photograph). *Cordylancistrus daguae*: ICNMHN 3515, 17643, 17644, 17645. *Cordylancistrus perijae*: ANSP 168917 (paratype), ICNMHN 17502 (ex CAR 270, ex MBUCV-V 21745, paratype). *Cordylancistrus platycephalus*: BMNH 1898.11.4:42 (holotype, photograph). *Cordylancistrus tayrona*: ICNMHN 17503 (ex CAR 370), MLS 541. *Cordylancistrus* sp2. "Pacífico": BMNH 1908.5.29.70-79. *Cordylancistrus* sp3. "Magdalena": CP-UCO 1060, 1062. *Cordylancistrus* sp4.: FMNH 76213 (cs). *Cordylancistrus torbesensis*: USNM 121001 (holotype, xray). *Dekeyseria niveata*: ANSP 185259. *Dekeyseria pulcher*: ANSP 185298. *Dekeyseria scaphirhyncha*: ICNMHN 12787, 12788. *Dolichancistrus atratoensis*: CIUA 768, 769, 771, 772, IAvH-P 6630, ICNMHN 51 (holotype), 46 (paratypes), 74 (paratype), 3460. *Dolichancistrus carnegiei*: ICNMHN 591, 1822, 3235, 3571, 5445, 16016, 16017, 16018, 17498, 17500, 17501, MLS 522,

542, 543, 550. *Dolichancistrus cobrensis*: AUM 30377, 46306, MCNG 541, USNM 121036 (holotype), 121037 (paratypes). *Dolichancistrus fuesslii*: IAvH-P 7931, 11381, 3939, 3940, 9230, 9605, ICNMHN 2638, 2817, 2839, 3212, 3641, 14582, 16811, NMW 48026. *Hemiancistrus sabaji*: ANSP 185153. *Hemiancistrus guahiborum*: ICNMHN 5323, 11915. *Hemiancistrus punctulatus*: ANSP 170168. *Hopliancistrus tricornis*: AUM 39853. *Hypancistrus contradens*: ICNMHN 11917, 11918. *Hypancistrus debilittera*: ICNMHN 10691. *Hypostomus* cf. *wachereri*: MZUSP 87480. *Lasiancistrus caucanus*: ICNMHN 8763. *Lasiancistrus guacharote*: ICNMHN 16916. *Lasiancistrus triactis*: ZMA 120774. *Leporacanthicus galaxias*: AUM 42144. *Leptoancistrus canensis*: USNM 78300 (paratypes, xray). *Leptoancistrus* cf. *cordobensis*: CIUA 774, 775, 776, 777, 778, 779, 780, 781. *Lithoxus jantjae*: ANSP 182809 (paratypes). *Lithoxus lithoides*: ANSP 39121 (paratype). *Megalancistus aculeatus*: USNM 52594. *Neblinichthys pilosus*: ANSP 157587 (paratypes). *Neblinichthys roraima*: ANSP 174914 (paratypes). *Panaque cochliodon*: ICNMHN 369. *Panaque maccus*: ICNMHN 15728. *Peckoltia bachi*: ICNMHN 13952. *Peckoltia brevis*: ICNMHN 7952. *Peckoltia vittata*: ICNMHN 7954, 12792. *Pseudacanthicus leopardus*: AUM 35550, USNM 197105. *Pseudacanthicus spinosus*: USNM 52594. *Pseudancistrus sidereus*: ANSP 185297. *Pseudolithoxus dumus*: ANSP 185255. *Spectracanthicus punctatissimus*: MNHN 1999-0021.

**MALAPTERURIDAE**

*Malapterurus beninensis*: MZUSP 84464, USNM 303492.

**MOCHOCKIDAE**

*Chiloglanis swierstrai*: MZUSP 65801. *Synodontis schall*: MZUSP 84468.

**NEMATOGENYIDAE**

*Nematogenys inermis*: MZUSP 75256.

**PANGASIIDAE**

*Pangasius macronema*: MZUSP 62606. *Pangasius pangasius*: UMMZ 208434.

**PIMELODIDAE**

*Bergiaria westermanni*: MZUSP 85627. *Brachyplatystoma capapretum*: MZUSP 53262, ANSP 178524, 179733, 178524. *Brachyplatystoma juruense*: ANSP 178514-1,3,4,6, DUF 1071. *Brachyplatystoma platynemum*: DUF 993, 1076, ANSP uncat, 187321. *Brachyplatystoma rousseauxii*: ANSP 179794, 179793, 179233, DUF 1051, 981, 1078. *Brachyplatystoma vaillanti*: ANSP 179799, 178525, 179474, DUF 994, 1164. *Brachyplaystoma filamentosum*: DUF 1079, 1080, ANSP 179776. *Calophysus macropterus*: DUF 1199, 1049, 403, ANSP 199813, 178164, 178260, ICNMHN uncat. *Duopalatinus emarginatus*: MZUSP 85622. *Exallodontus aguanai*: ANSP 18947, uncat. *Hemisorubim platyrhynchos*: ANSP uncat, 179234, ICNMHN 7909, MZUSP 7009. *Hypophthalmus* sp. “curved”: ANSP uncat. *Hypophthalmus* sp. “straight”: ANSP 180993, 178512, 180993, 187103. *Iheringichthys labrosus*: ANSP 180505, MZUSP 78459, 25102. *Leiarius marmoratus*: MZUSP 108333. *Leiarius perruno*: ANSP uncat-I, uncat-II, ICNMHN 2158. *Leiarius pictus*: MZUSP 82577. *Leiarius* sp.: ANSP 178526, 178527, uncat without skull, DUF 1054, 1056, 1036, 1055, 1037. *Luciopimelodus pati*: ANSP 178798, MZUSP 78464, 78457. *Megalonema platanum*: MZUSP 78465. *Megalonema platycephalum*: ANSP 179249, 178515. *Megalonema* sp.: MZUSP 92604. *Parapimelodus nigribarbis*: MZUSP 78451. *Parapimelodus valenciennis*: MZUSP 78466, ANSP 178800. *Phractocephalus hemioliopterus*: ANSP 179559, 179553, 179554, ICN uncat. *Pimelodina flavipinnis*: ANSP uncat, 178513, 178516. *Pimelodus argenteus*: ANSP 181017, uncat. *Pimelodus fur*: MZUSP 22566. *Pimelodus grosskopfii*: ICNMHN 6867.

*Pimelodus maculatus*: MZUSP 85486, 110379. *Pimelodus microstoma*: MZUSP 22696, 22712 (paratype of *Pimelodus heraldoi*). *Pimelodus mysteriosus*: ANSP 180506, MZUSP 90595. *Pimelodus ortmanni*: MZUSP 50053. *"Pimelodus" ornatus*: ANSP 178452, 180985, MZUSP 34480, 109128. *"Pimelodus"* sp.: MZUSP 58328. *Pinirampus argentina*: ANSP 181016. *Pinirampus pirinampu*: ANSP uncat, 178530. *Platynematichthys notatus*: ANSP uncat, 178528. *Platysilurus malarmo*: ANSP uncat, IAvH-P 11815, IAvH-P 11819, IAvH-P 15894. *Platysilurus mucosus*: DUF 986, uncat, ANSP 178508, ANSP 178509, IAvH-P 17638, IAvH-P 10850, IAvH-P 10856, IAvH-P 6006. *Platystomatichthys sturio*: ANSP uncat, SU 22463, ICNMHN 15201. *Platystomatichthys* sp. "Xingu": MZUSP 58364. *Propimelodus* sp.: ANSP 180939. *Pseudoplatystoma corruscans*: ANS 188913, MZUSP 78477. *Pseudoplatystoma fasciatum*: ANSP 177346. *Pseudoplatystoma magdaleniatum*: ICNMHN 6860. *Pseudoplatystoma metense*: ANSP 149541. *Pseudoplatystoma reticulatum*: ANSP 188912. *Pseudoplatystoma* sp.: DUF 1125, ICNMHN uncat. *Pseudoplatystoma tigrinum*: DUF 921, uncat, ANSP 187010. *Sorubim cuspicaudus*: MBUCV uncat, DUF 932. *Sorubim elongatus*: ICNMHN 15039. *Sorubim lima*: ANSP 178507. *Sorubim trigonocephalus*: ANSP 188824. *Sorubimichthys planiceps*: 179235, 17850. *Steindachneridion scripta*: MZUSP 78463. *Zungaro zungaro*: ANSP uncat, DUF 982, ICN uncat, MPUJ 13213, MZUSP 96290, 108335, 94859, 96188, 151727.

**PLOTOSIDAE**

*Plotosus lineatus*: MZUSP 22179.

**PSEUDOPIMELODIDAE**

*Batrochoglanis villosus*: MZUSP 7356, 23864. *Cephalosilurus apurensis*: DU-F 924, 980, 1102, 1040, ICNMHN uncat. *Cephalosilurus fowleri*: MZUSP 73756. *Pseudopimelodus mangurus*: MZUSP 40684.*Pseudopimelodus raninus*: ICNMHN 6865. *Pseudopimelodus* sp.: MZUSP 78458. *Pseudopimelodus bufonius*: ANSP uncat, MBUCV 2001-12.6.

**SCHILBEIDAE**

*Schilbe intermedius*: MZUSP 62627. *Schilbe* sp.: MZUSP 65820.

**SILURIDAE**

*Ompok pabo*: MZUSP 63582. *Wallago leerii*: MZUSP 63357.

**SISORIDAE**

*Bagarius* cf. *yarrelli*: MZUSP 63119. *Erethistes hara*: MZUSP 48644. *Gagata cenia*: MZUSP 42052. *Pseudecheneis sulcata*: MZUSP 50844.

**TRICHOMYCTERIDAE**

*Trichomycterus iheringi*: MZUSP 120974.

**DIPNOI**

**LEPIDOSIRENIDAE**

*Lepidosiren paradoxa*: MZUSP 35634, 41101, 50036, 35633.

### 2 Phylogenetic analysis

#### 2.1 Introduction

The present analysis puts the fossil occurrences of *Phractocephalus* from the Sincelejo Fm. in a phylogenetic context. We included the morphological characters from Aguilera et al. (2008)

and Azpelicueta and Cione (2016) along with our own characters relevant to the systematics of the genus, that includes a single extant, and three extinct species. We carried out a combined bayesian phylogenetic analysis placing the fossil taxa along with recent taxa, using both morphological and molecular data using bayesian phylogenetic inference.

#### 2.2 Morphological characters

Previous studies have used morphological characters for inferring the interrelationships among species of *Phractocephalus*. We extended the original matrix of Aguilera et al. (2008) including the recently-described *Phractocephalus ivy* (Azpelicueta and Cione, 2016) and the specimens from the Sincelejo formation in order to assess their phylogenetic placement (Table 1). Characters were coded for *Phractocephalus acreornatus*, *P. ivy*, and *P. nassi* following Aguilera et al. (2008), Azpelicueta and Cione (2016), and Lundberg and Aguilera (2003) respectively. Extant species were coded examining osteological specimens. We included two additional new characters to this matrix.

##### 2.2.1 Characters

Characters 1 to 6 were taken from Aguilera et al. (2008) with the inclusion of a new one (character 7) herein proposed. An additional character was added from recent examination of cranial osteology in the Pimelodidae (character 8). Character definitions and states are described below.

**Character 1:** Skull ornamentation near midline. **States:** Weakly or strongly ridged and grooved (0). Mostly reticulating ridges and pits (1). **Comments:** *Phractocephalus ivy* was coded from figure 3 in Azpelicueta and Cione (2016). Impossible to code in our specimens of *Phractocephalus* Corozal.

**Character 2:** Mesethmoid width. **States:** Narrow (0). Very broad (1). **Comments:** *Phractocephalus ivy* was coded as missing (?) given that the character is not preserved in the available specimens of the original description. Impossible to code in our specimens of *Phractocephalus* Corozal.

**Character 3:** Anterior and lateral expansions of lateral ethmoid. **States:** Expansions absent (0). Expansions present but weak (1). Expansions present and strong (2). **Comments:** Said to be the same condition in *P. ivy* as in *Phractocephalus hemioliopterus* (Azpelicueta and Cione, 2016, p. 255). Impossible to code in our specimens of *Phractocephalus* Corozal.

**Character 4:** Length of supraoccipital process as compared to the length of Weberian apparatus. **States:** Supraoccipital process shorter than Weberian process (0). Supraoccipital process longer than Weberian process (1). **Comments:** Coded as missing (?) for *P. ivy* given that the character is not preserved in the available specimens of the original description. This character was coded as state 2 for *P. acreornatus* and *P. nassi* (Aguilera et al., 2008, table 2); however, the character definition only includes two states, 0 and 1. This coding error was corrected in the present matrix where the species in question should have state 1 instead. Impossible to code in our specimens of *Phractocephalus* Corozal.

**Character 5:** Degree of development of ornamentation on opercle. **States:** Localized and poorly-developed (0). Covering most of the opercle but moderately-developed (1). Covering most of the opercle and strongly-developed (2). **Comments:** Said to be the same condition in *P. ivy* as in *P. nassi* and *P. acreornatus* (Azpelicueta and Cione, 2016, p. 255). Impossible to code in our specimens of *Phractocephalus* Corozal.

**Character 6:** Ornamentation on the pectoral-spine shaft. **States:** Consisting of semi-parallel ridges or shaft surface smooth (0). Consisting of bony reticulations and some pits inbetween (1). Consisting of strong reticulations that form extensive cover of pits on the spine shaft (2). **Comments:** Redefined in the present work and coded from the figure 9 in Azpelicueta and Cione (2016) for *P. ivy*.

**Character 7:** Bifid spinules on proximal section of posterior pectoral-spine shaft. **States:** Absent (0). Present (1). **Comments:** New character, coded from figure 9 of (Azpelicueta and Cione, 2016) for *P. ivy*, from figure 5 and explicitly from text (p.104) in Lundberg and Aguilera (2003) for *P. nassi*, and from figure 10 in Aguilera et al. (2008) for *P. acreornatus*.

**Character 8:** Contribution of the lateral ethmoid to the margin of the orbit. **States:** Less than half of the dorsal margin of the orbit (0). More than half of the dorsal margin of the orbit (1). **Comments:** New character, coded from specimens examined in extant species, and from figure 7 of Azpelicueta and Cione (2016) for *P. ivy*, from figure 3 of Lundberg and Aguilera (2003) for *P. nassi*, and from figure 2 of Aguilera et al. (2008) for *P. acreornatus*. Impossible to code in our specimens of *Phractocephalus* Corozal.

##### 2.2.2 Morphological matrix (modified from Aguilera et al. (2008))

The modified matrix is herein presented with codes indicating the specific source of coding.

Citation codes for the table:

- `AC`: Azpelicueta and Cione (2016)
- `AE`: Aguilera et al. (2008)
- `LA`: Lundberg and Aguilera (2003)

Table 1: Morphological characters used in the phylogenetic analysis. Missing data (?) are indicated whenever the condition could not be unambiguously coded. The codes by each character state indicate the figure `f` or page `p` where the state can be evidenced in the relevant references.

| Taxon and characters | Ch1 | Ch2 | Ch3 | Ch4 | Ch5 | Ch6 | Ch7 | Ch8 |
|---|---|---|---|---|---|---|---|---|
| Outgroup (composite) | 0 | 0 | 0 | 0 | 0 | 0 | 0 | 0 |
| *P. acreornatus* | 0 | 1 | 2 | 1 | 2 | 2 | 1 `AEf10` | 1 `AEf2` |
| *P. ivy* | 0 `ACf3` | ? | 0 `ACp255` | ? | 2 `ACp255` | 1 `ACf9` | 1 `ACf9` | 1 `ACf7` |
| *P. nassi* | 0 | 1 | 1 | 1 | 2 | 1 | 1 `LAf5p104` | 1 `LAf3` |
| *P. hemioliopterus* | 1 | 0 | 0 | 1 | 1 | 2 | 1 | 1 |
| *Phractocephalus* Corozal | ? | ? | ? | ? | ? | 1 | 1 | ? |

Please note that while Aguilera et al. (2008) treated the outgroup as a truly composite entry in the matrix for inference, our current instance is composite only for brevity as it is invariant in all the characters presented herein, it will be coded accordingly when concatenated to the molecular partitions.

##### 2.2.3 Dataset in nexus format

This preamble is important (special parts in letters, e.g., A, B, etc.)

```
#NEXUS
BEGIN DATA;
[number of taxa in ntax and number of characters (total) in nchar]
dimensions ntax=84 nchar=7578;
[Standard are the morpho chars and DNA the molec chars, they must start in
7571 and end in 7578 position (morpho) and start in 1 and end in 7570 (molec)]
['interleave' indicates that the matrix can be specified in blocks in order
to avoid pasting all characters in a single string]
format datatype=mixed(DNA: 1-7570, Standard: 7571-7578) gap=- missing=? interleave=yes;
```

When specifying the partitions by position (the `charset` command) use the following commands. This will count the partitions (Nparts = morphology + molecular partitions) and define set them to be combined with the partition name used below in `charset`. unlink parameters is important in order to allow them to be governed by their own parameters during likelihood calculations:

```
CHARSET morphology = A-B;
...

partition combined = Nparts :  morphology, molecular_partitions;
set partition = combined;
unlink shape = ( all ) pinvar = ( all ) statefreq = ( all ) revmat = ( all );
```

Also, it is important to allow al partitions to have their own rates, so the next line is important:

```
[allow separate gamma parameters for each partition]
prset applyto=(all) ratepr=variable;
```

##### 2.2.4 Morphological substitution model

There are two substitution models used in MrBayes 3: Lewis 2001 and parsimony. The default is Lewis (2001) that is analog to JC except for a variable number of states, up to 10. On the other hand, the "parsimony" model is much parameter-rich than other models and therefore is not parsimonious in statistical terms given that each character will require its own parameter set, and therefore the whole model will have $r * n$ parameters where $r$ is the number of parameter for each character, and $n$ is the number of characters. Therefore, the best agnostic model would be Lewis (2001) where the whole morphological partition is governed by the same set of parameters.

One important aspect of determining the morphological partition and specifying its substitution model is to define the sampling bias, that is, how did we sample the morphological characters. If we sampled continuously so that theoretically every morphological character had the same probability of being sampled, the parameter `coding` should take the value `all`. On the other hand, if we sampled so that characters that show only 0 or 1 in all taxa can not be observed, the value should be `variable`. Finally, if we sampled only parsimony-informative characters, the value should be set to `informative`. This is really important in model-based approaches where the frequencies of states are expected to inform about the real frequencies and therefore guide the model updating during MCMC. This is called ascertainment bias when morphology is sampled in the classical way. There seems to be no much to do besides specifying either `variable` or `informative`. The important difference here between the latter two values is that variable characters need not to be informative under parsimony. This is specified under the `lset` command:

```
[morphology substitution model]
lset applyto=(1) coding=variable;
```

There seems to be no need to specify the substitution model as the default (`Lewis 2001`) is enough for this partition.

#### 2.3 Taxon sets

The following taxa are considered in the analysis:

```
Brachyplatystoma_capapretum
Brachyplatystoma_filamentosum_179214
Brachyplatystoma_filamentosum_179446
Brachyplatystoma_filamentosum_187105
Brachyplatystoma_juruense_179217
Brachyplatystoma_juruense_179454
Brachyplatystoma_platynemum
Brachyplatystoma_rousseauxii_179233
Brachyplatystoma_vaillantii_178525
Brachyplatystoma_vaillantii_187107
Calophysus_macropterus
Cheirocerus_abuelo
Cheirocerus_eques
Duopalatinus_peruanus
Exallodontus_aguanai_179220
Exallodontus_aguanai_MBUCV_VCD_8
Goeldiella_eques
Hemisorubim_platyrhynchos
Hypophthalmus_cf_edentatus
Hypophthalmus_cf_marginatus
Hypophthalmus_fimbriatus
Iheringichthys_labrosus
Leiarius_marmoratus
Leiarius_pictus
Luciopimelodus_pati
Megalonema_amaxanthum
Megalonema_platanum
Megalonema_platycephalum_178450
Parapimelodus_nigribarbis
Parapimelodus_valenciennis
Perrunichthys_perruno
Phractocephalus_hemioliopterus
Pimelodidae_sp.1
Pimelodidae_sp.2
Pimelodidae_sp.3
Pimelodidae_sp.4
Pimelodina_flavipinnis
Pimelodus_albicans
Pimelodus_argenteus
Pimelodus_blochii_179455
Pimelodus_blochii_179456
Pimelodus_blochii_179457
Pimelodus_blochii_49101
Pimelodus_blochii_54570
Pimelodus_cf_blochii_179186
Pimelodus_cf_altissimus_179190
Pimelodus_cf_altissimus_179226
Pimelodus_coprophagus
Pimelodus_maculatus_181109
```

```
Pimelodus_maculatus_78452
Pimelodus_ornatus_178452
Pimelodus_ornatus_22394
Pimelodus_ornatus_49102
Pimelodus_ornatus_5131
Pimelodus_pictus_178168
Pimelodus_pictus_179448
Pinirampus_pirinampu_178265
Pinirampus_pirinampu_180992
Pinirampus_pirinampu_181016
Platynematichthys_notatus
Platysilurus_malarmo
Platysilurus_mucosus_179215
Platysilurus_mucosus_22654
Platystomatichthys_sturio
Pseudopimelodus_mangurus
Pseudoplatystoma_corruscans
Pseudoplatystoma_fasciatum
Pseudoplatystoma_magdaleniatum
Pseudoplatystoma_tigrinum
Sorubim_cuspicaudus
Sorubim_elongatus
Sorubim_lima
Sorubim_maniradii
Sorubimichthys_planiceps
Steindachneridion_scriptum
Zungaro_zungaro
Brachyplatystoma_rousseauxii_187109
Brachyplatystoma_tigrinum
Megalonema_platycephalum_179458
Leiarius_longibarbis
Steindachneridion_scripta
Hypophthalmus_edentatus
```

Some taxa were unavailable for analysis due to the lack of osteological specimens for examination. These are *Brachyplatystoma tigrinum*, *Cheirocerus abuelo*, *Cheirocerus eques*, *Hypophthalmus fimbriatus*, *Leiarius longibarbis*, *Megalonema amaxanthum*, *Pimelodus albicans*, *Pimelodus* cf. *altissimus*, *Pimelodus coprophagus*, *Pimelodus pictus*, and *Sorubim maniradii.* However, there were molecular data for these and consequently they were included in the analysis applying missing data to the morphological partition in their respective positions.

Note: This list was produced by removing whatever between blank spaces and end of line with the regexp `\s-\(.*\)$` with the keystroke `C-M-%` (or `Ctrl+Alt+Shift+5`) for `replace-regexp` in Emacs.

#### 2.4 Molecular data

Fossil terminals with missing data for molecular partitions were filled with `(make-string N ??)` where N is the number of missing data in a row, and `??` represents "?" in the Emacs function `make-string`. Given that *Phractocephalus hemioliopterus* is already represented in the molecular alignment, only the three extinct species and the Corozal specimen needed to be placed after all other taxa in the molecular matrix:

```
...
Steindachneridion_scripta          ---------------------------------
Hypophthalmus_edentatus            ---------------------------------
Phractocephalus_acreornatus        ?????????????????????????????????
Phractocephalus_ivy                ?????????????????????????????????
Phractocephalus_nassi              ?????????????????????????????????
Phractocephalus_Corozal            ?????????????????????????????????
...
```

#### 2.5 Analysis implementation

The dataset (in nexus format) and scripts (in `R`, `bash`, and `MrBayes`) for reproducing the analysis are available in THE DIRECTORY xxxxx.

#### 2.6 Analysis results

The following is an excerpt of relevant sections of the `.log` file:

```
...
Average standard deviation of split frequencies: 0.004698

Analysis completed in 4 hours 17 mins 20 seconds
Analysis used 15433.77 seconds of CPU time on processor 0
Likelihood of best state for "cold" chain of run 1 was -61192.14
Likelihood of best state for "cold" chain of run 2 was -61196.45
...
Summary statistics for partitions with frequency >= 0.10 in at least one run:
    Average standard deviation of split frequencies = 0.004698
    Maximum standard deviation of split frequencies = 0.032954
    Average PSRF for parameter values (excluding NA and >10.0) = 1.000
    Maximum PSRF for parameter values = 1.008
...
# 98% HPD interval remomved from this excerpt for brevity...
Parameter       Mean      Variance     Median    min ESS*  avg ESS    PSRF+
-------------------------------------------------------------------------
TL{all}       2.400185    0.009124    2.395997    626.76    688.88    1.002
kappa{2}      3.989707    0.105469    3.975593    458.80    526.19    1.000
kappa{4}     12.798448    0.379460   12.784600    621.71    631.22    0.999
r(A<->C){1}   0.089537    0.000053    0.089183    671.20    711.10    0.999
r(A<->G){1}   0.280846    0.000213    0.280674    648.14    670.27    1.000
r(A<->T){1}   0.085786    0.000054    0.085905    554.95    652.97    1.001
r(C<->G){1}   0.065785    0.000042    0.065631    751.00    751.00    1.000
r(C<->T){1}   0.419569    0.000282    0.419714    628.79    683.99    1.001
r(G<->T){1}   0.058478    0.000038    0.058417    751.00    751.00    0.999
r(A<->C){3}   0.088174    0.000027    0.088035    603.87    677.44    1.000
r(A<->G){3}   0.222331    0.000183    0.222310    336.91    422.57    1.000
r(A<->T){3}   0.064775    0.000023    0.064663    682.01    695.35    1.000
r(C<->G){3}   0.005524    0.000004    0.005292    621.49    686.25    1.004
r(C<->T){3}   0.603542    0.000234    0.603397    484.89    527.91    1.001
r(G<->T){3}   0.015654    0.000009    0.015397    633.68    692.34    0.999
pi(A){1}      0.290219    0.000065    0.290229    663.63    683.70    1.001
pi(C){1}      0.221970    0.000050    0.221863    565.40    658.20    1.000
pi(G){1}      0.238408    0.000055    0.238417    704.18    727.59    0.999
pi(T){1}      0.249403    0.000057    0.249463    609.71    680.35    0.999
pi(A){2}      0.268564    0.000126    0.268310    751.00    751.00    0.999
pi(C){2}      0.233198    0.000124    0.233258    751.00    751.00    0.999
pi(G){2}      0.233218    0.000122    0.233088    649.40    700.20    0.999
pi(T){2}      0.265019    0.000118    0.265256    710.85    730.93    1.000
pi(A){3}      0.363814    0.000068    0.363738    695.39    723.20    1.002
pi(C){3}      0.237466    0.000046    0.237286    549.20    605.30    1.000
pi(G){3}      0.185158    0.000053    0.185276    579.42    611.87    1.001
pi(T){3}      0.213562    0.000042    0.213632    570.11    623.77    0.999
pi(A){4}      0.335488    0.000091    0.335634    751.00    751.00    0.999
pi(C){4}      0.367595    0.000072    0.367705    669.46    710.23    0.999
pi(G){4}      0.083074    0.000012    0.082991    751.00    751.00    1.000
pi(T){4}      0.213843    0.000036    0.213932    682.69    716.85    0.999
alpha{1}      0.929507    0.021963    0.916908    651.90    701.45    1.001
alpha{2}      0.845560    0.039927    0.824301    665.83    708.41    1.000
alpha{3}      0.628006    0.004524    0.624135    643.79    693.45    1.001
alpha{4}      0.858176    0.003148    0.856519    751.00    751.00    0.999
pinvar{1}     0.393908    0.001537    0.396444    611.27    661.28    1.000
pinvar{2}     0.296148    0.004579    0.302666    574.05    628.66    1.003
pinvar{3}     0.558731    0.000429    0.559689    648.29    653.72    1.000
pinvar{4}     0.479202    0.000267    0.479482    751.00    751.00    1.000
```

```
m{1}         0.344125    0.000267    0.343594    751.00    751.00    1.001
m{2}         0.437752    0.000812    0.436977    628.68    656.28    1.000
m{3}         0.885944    0.001246    0.886473    581.53    593.41    1.000
m{4}         3.066338    0.009307    3.066057    569.30    660.15    1.000
m{5}         0.545579    0.063024    0.516275    616.93    657.73    0.999
------------------------------------------------------------------------
* Convergence diagnostic (ESS = Estimated Sample Size); min and avg values
  correspond to minimal and average ESS among runs.
  ESS value below 100 may indicate that the parameter is undersampled.
+ Convergence diagnostic (PSRF = Potential Scale Reduction Factor; Gelman
  and Rubin, 1992) should approach 1.0 as runs converge.
```

#### 2.7 Accession numbers

| **Catalog Number** | **Species** | ***cytb*** | ***rag1*** | ***rag2*** | ***12S*** |
|---|---|---|---|---|---|
| ANSP 179218 | *Brachyplatystoma capapretum* | JF898520 | JF898598 | JF898745 | JF898674 |
| ANSP 179214 | *Brachyplatystoma filamentosum* | JF898521 | JF898599 | JF898746 | JF898675 |
| ANSP 179446 | *Brachyplatystoma filamentosum* | JF898522 | JF898600 | JF898747 | JF898676 |
| ANSP 187105 | *Brachyplatystoma filamentosum* | JF898523 | JF898601 | JF898748 | JF898677 |
| ANSP 179217 | *Brachyplatystoma juruense* | JF898516 | JF898594 | JF898741 | JF898670 |
| ANSP 179454 | *Brachyplatystoma juruense* | JF898517 | JF898595 | JF898742 | JF898671 |
| ANSP 179230 | *Brachyplatystoma platynemum* | JF898515 | JF898593 | JF898740 | JF898669 |
| ANSP 179233 | *Brachyplatystoma rousseauxii* | JF898518 | JF898596 | JF898743 | JF898672 |
| ANSP 187109 | *Brachyplatystoma rousseauxii* | JF898519 | - | JF898744 | JF898673 |
| ANSP 179236 | *Brachyplatystoma tigrinum* | JF898514 | - | JF898739 | JF898668 |
| ANSP 178525 | *Brachyplatystoma vaillantii* | JF898512 | JF898590 | JF898737 | JF898666 |
| ANSP 187107 | *Brachyplatystoma vaillantii* | JF898513 | JF898591 | JF898738 | JF898667 |
| ANSP 179229 | *Calophysus macropterus* | JF898528 | JF898607 | JF898753 | JF898682 |
| MBUCV 2001.12.6.L03 | *Cheirocerus abuelo* | JF898533 | JF898612 | JF898758 | JF898687 |
| INHS 52717 | *Cheirocerus eques* | JF898534 | JF898613 | JF898759 | JF898688 |
| ANSP 179221 | *Duopalatinus peruanus* | JF898539 | JF898618 | JF898763 | JF898693 |
| ANSP 179220 | *Exallodontus aguanai* | JF898542 | JF898621 | JF898765 | JF898696 |
| MBUCV-VCD#8 | *Exallodontus aguanai* | JF898543 | JF898622 | JF898766 | JF898697 |
| INHS 49299 | *Goeldiella eques* | JF898565 | JF898644 | DQ492368 | JF898719 |
| ANSP 179234 | *Hemisorubim platyrhynchos* | - | JF898588 | JF898735 | JF898664 |
| INHS 52182 (cytb, rag1, 12S) / INHS 5218 (rag2) | *Hypophthalmus* cf. *edentatus* | JF898505 | JF898583 | DQ492362 | JF898661 |
| ANSP 179219 | *Hypophthalmus* cf. *marginatus* | JF898504 | JF898582 | JF898730 | JF898660 |
| ANSP 179185 | *Hypophthalmus fimbriatus* | - | JF898581 | JF898729 | JF898659 |
| ANSP 180505 | *Iheringichthys labrosus* | JF898550 | JF898629 | JF898773 | JF898704 |
| ANSP 178109 | *Leiarius longibarbis* | DQ486766 | - | DQ486784 | - |
| ANSP 178109 | *Leiarius marmoratus* | - | JF898568 | - | JF898647 |
| INHS 54805 | *Leiarius perruno* | DQ486767 | JF898570 | DQ486785 | - |
| ANSP 178108 | *Leiarius pictus* | DQ486764 | JF898569 | DQ486782 | JF898648 |
| MZUSP 78457 | *Luciopimelodus pati* | JF898529 | JF898608 | JF898754 | JF898683 |
| ANSP 179213 | *Megalonema amaxanthum* | JF898535 | JF898614 | - | JF898689 |
| MZUSP 78465 | *Megalonema platanum* | JF898536 | JF898615 | JF898760 | JF898690 |
| ANSP 179458 | *Megalonema platycephalum* | JF898537 | - | JF898761 | JF898691 |
| ANSP 178450 | *Megalonema platycephalum* | JF898538 | JF898617 | JF898762 | JF898692 |
| MZUSP 78451 | *Parapimelodus nigribarbis* | JF898551 | JF898630 | JF898774 | JF898705 |

| Catalog Number | Species | cytb | rag1 | rag2 | 12S |
|---|---|---|---|---|---|
| MZUSP 78466 | *Parapimelodus valenciennis* | JF898552 | JF898631 | - | JF898706 |
| ANSP 179554 (cytb, rag1, 12S) / ANSP 179452 (rag2) | *Phractocephalus hemioliopterus* | JF898492 | JF898567 | DQ486781 | JF898646 |
| ANSP 179223 | Pimelodidae sp. 1 | - | JF898623 | JF898767 | JF898698 |
| ANSP 179231 | Pimelodidae sp. 2 | JF898545 | JF898624 | JF898768 | JF898699 |
| ANSP 185247 | Pimelodidae sp. 3 | JF898546 | JF898625 | JF898769 | JF898700 |
| ANSP 179187 | Pimelodidae sp. 4 | JF898547 | JF898626 | JF898770 | JF898701 |
| ANSP 179225 | *Pimelodina flavipinnis* | JF898527 | JF898606 | JF898752 | JF898681 |
| ANSP 178802 | *Pimelodus albicans* | JF898553 | JF898632 | JF898776 | JF898707 |
| ANSP 181017 | *Pimelodus argenteus* | JF898557 | JF898636 | JF898780 | JF898711 |
| INHS 49101 | *Pimelodus blochii* | JF898559 | JF898638 | JF898782 | JF898713 |
| ANSP 179455 | *Pimelodus blochii* | JF898560 | JF898639 | JF898783 | JF898714 |
| INHS 54570 | *Pimelodus blochii* | JF898561 | JF898640 | JF898784 | JF898715 |
| ANSP 179456 | *Pimelodus blochii* | JF898562 | JF898641 | JF898785 | JF898716 |
| ANSP 179457 | *Pimelodus blochii* | JF898563 | JF898642 | JF898786 | JF898717 |
| ANSP 179190 | *Pimelodus* cf. *altissimus* | JF898540 | JF898619 | - | JF898694 |
| ANSP 179226 | *Pimelodus* cf. *altissimus* | JF898541 | JF898620 | JF898764 | JF898695 |
| ANSP 179186 | *Pimelodus* cf. *blochii* | JF898558 | JF898637 | JF898781 | JF898712 |
| MBUCV 2001.12.6.L10 | *Pimelodus coprophagus* | JF898556 | JF898635 | JF898779 | JF898710 |
| ANSP 181109 | *Pimelodus maculatus* | JF898554 | JF898633 | JF898777 | JF898708 |
| MZUSP 78452 | *Pimelodus maculatus* | JF898555 | JF898634 | JF898778 | JF898709 |
| INHS 49102 | *Pimelodus ornatus* | JF898524 | JF898602 | DQ492363 | JF898678 |
| ANSP 178452 | *Pimelodus ornatus* | JF898525 | JF898603 | JF898749 | JF898679 |
| AUM 22394 | *Pimelodus ornatus* | JF898526 | JF898604 | JF898750 | - |
| MLP 5131 | *Pimelodus ornatus* | - | JF898605 | JF898751 | JF898680 |
| ANSP 178168 | *Pimelodus pictus* | JF898548 | JF898627 | JF898771 | JF898702 |
| ANSP 179448 | *Pimelodus pictus* | JF898549 | JF898628 | JF898772 | JF898703 |
| ANSP 178265 | *Pinirampus pirinampu* | JF898530 | JF898609 | JF898755 | JF898684 |
| ANSP 180992 | *Pinirampus pirinampu* | JF898531 | JF898610 | JF898756 | JF898685 |
| ANSP 181016 | *Pinirampus pirinampu* | JF898532 | JF898611 | JF898757 | JF898686 |
| ANSP 179227 | *Platynematichthys notatus* | JF898511 | JF898589 | JF898736 | JF898665 |
| MBUCV 2001.12.6.L17 | *Platysilurus malarmo* | JF898507 | JF898585 | JF898732 | JF898662 |
| ANSP 179215 | *Platysilurus mucosus* | JF898508 | JF898586 | JF898733 | - |
| AUM 22654 | *Platysilurus mucosus* | JF898509 | JF898587 | JF898734 | JF898663 |
| ANSP 179216 | *Platystomatichthys sturio* | JF898506 | JF898584 | JF898731 | - |
| MZUSP 78458 | *Pseudopimelodus mangurus* | JF898564 | JF898643 | DQ492360 | JF898718 |
| MZUSP uncat. skel | *Pseudoplatystoma corruscans* | JF898500 | JF898578 | JF898727 | JF898657 |
| ANSP 179449 | *Pseudoplatystoma fasciatum* | JF898501 | JF898579 | - | - |
| ANSP 188882 | *Pseudoplatystoma magdaleniatum* | JF898499 | JF898577 | JF898726 | JF898656 |
| ANSP 179450 | *Pseudoplatystoma tigrinum* | JF898502 | JF898580 | JF898728 | JF898658 |
| AUM 22581 | *Sorubim cuspicaudus* | JF898494 | JF898572 | JF898721 | JF898651 |
| ANSP 178315 | *Sorubim elongatus* | JF898497 | JF898575 | JF898724 | JF898654 |
| ANSP 178507 | *Sorubim lima* | JF898496 | JF898574 | JF898723 | JF898653 |
| ANSP 178366 | *Sorubim maniradii* | JF898495 | JF898573 | JF898722 | JF898652 |
| INHS 54701 | *Sorubimichthys planiceps* | JF898498 | JF898576 | JF898725 | JF898655 |
| MZUSP 78463 | *Steindachneridion scriptum* | DQ486765 | JF898566 | DQ486783 | JF898645 |
| INHS 43365 | *Zungaro zungaro* | JF898493 | JF898571 | JF898720 | JF898650 |
